# Spatial transcriptomics of a parasitic flatworm provides a molecular map of vaccine candidates, drug targets and drug resistance genes

**DOI:** 10.1101/2023.12.11.571084

**Authors:** Svenja Gramberg, Oliver Puckelwaldt, Tobias Schmitt, Zhigang Lu, Simone Haeberlein

## Abstract

The spatial organization of gene expression dictates tissue functions in multicellular parasites. Here, we present the first spatial transcriptome of a parasitic flatworm, the common liver fluke *Fasciola hepatica.* We identified gene expression profiles and marker genes for eight distinct tissues and validated the latter by *in situ* hybridization. To demonstrate the power of our spatial atlas, we focused on genes with substantial medical importance, including vaccine candidates (Ly6 proteins), drug targets (β-tubulins, protein kinases) and drug resistance genes (glutathione S-transferases, ABC transporters). Several of these genes exhibited unique expression patterns, indicating tissue-specific biological functions. Notably, the prioritization of tegumental protein kinases identified a PKCβ, for which small-molecule targeting caused parasite death. Our comprehensive gene expression map provides unprecedented molecular insights into the organ systems of this complex parasitic organism, serving as a valuable tool for both basic and applied research.

## Introduction

Latest developments in "omics” technologies have revolutionized biomedical research in recent years, with notable contributions from single-cell transcriptomics and more recently, spatial transcriptomics. While single-cell transcriptomics is able to provide highly resolved information on cell-type specific gene expression, a drawback is the lack of spatial information that may be crucial to understand gene function. This limitation is complemented by spatial transcriptomics, which provides transcriptome-wide and spatially resolved gene expression data^1^. The ability to link transcriptomic profiles to anatomical features of tissues has proven invaluable in elucidating disease processes, especially in the fields of oncology, neurology and immunology^2–4^. Additionally, spatial transcriptomics is equally valuable for enhancing our understanding of the biology of entire complex organisms, including their organ function and development. Pioneering work in this direction has recently been made for model organisms such as *Drosophila* sp. and planarians^5–7^. With our work, we want to highlight that this cutting-edge technology should not be reserved for model organisms and human research, but can also improve our understanding of the biology of multicellular pathogens in an unprecedented way. In this spirit, our study represents the first application of spatially resolved transcriptomics to a parasitic flatworm, the common liver fluke *Fasciola hepatica*.

*F. hepatica*, together with related species, is the causative agent of fascioliasis, a zoonotic disease affecting at least 2.4 million people and numerous livestock worldwide^8^. Adult parasites are large multicellular organisms residing in the bile ducts of the host’s liver. They are dorso-ventrally flattened, leaf-like in shape and composed of a skin-like tegument, muscular suckers, a branched intestine, complex reproductive organs and further, largely uncharacterized tissues^9^.

To date, triclabendazole (TCBZ) is the only drug that is effective against almost all intra-mammalian life stages of liver flukes, but reports of TCBZ-resistant parasite strains are increasing^10,11^. This drives global research endeavors to find alternative treatments and vaccines^12,13^. A key assumption in target-based compound screenings is that compounds or their derivatives that interfere with the function of an essential molecule/protein will cause death of the parasite^14^. To this end, it seems more promising to target proteins expressed in organs essential for liver fluke survival, such as the tegument or the intestine. A spatial transcriptome would provide the information needed to classify and prioritize potential drug and vaccine targets based on their spatial expression in different organs.

Therefore, we intended to generate a dataset facilitating the rapid and uncomplicated evaluation of the spatial expression of thousands of liver fluke genes. This resource will enhance our comprehension of tissue-associated gene function and thereby help us to gain new insights into the molecular cell biology of this complex parasitic organism. With this work, we demonstrate the capability of spatial transcriptomics in advancing parasite research, serving as a source of inspiration for both fundamental questions and the development of new drugs and vaccines.

## Results

### Identification of eight transcriptionally distinct tissues in adult liver flukes

Employing the 10x Genomics Visium technology, we constructed a transcriptomic map of the adult stage of *F. hepatica*, the life stage causing chronic liver disease. In order to achieve maximum release of high-quality RNA from cryosections of the parasite, we first optimized the tissue permeabilization time using the 10x Visium tissue optimization workflow. Subsequently, we processed four transversal cryosections, each containing a different set of tissues (Figure S1a), using the 10x Visium spatial gene expression platform and Illumina sequencing to obtain spatially resolved gene expression data from those sections. An overview of the workflow is shown in Figure 1a.

**Figure 1.**
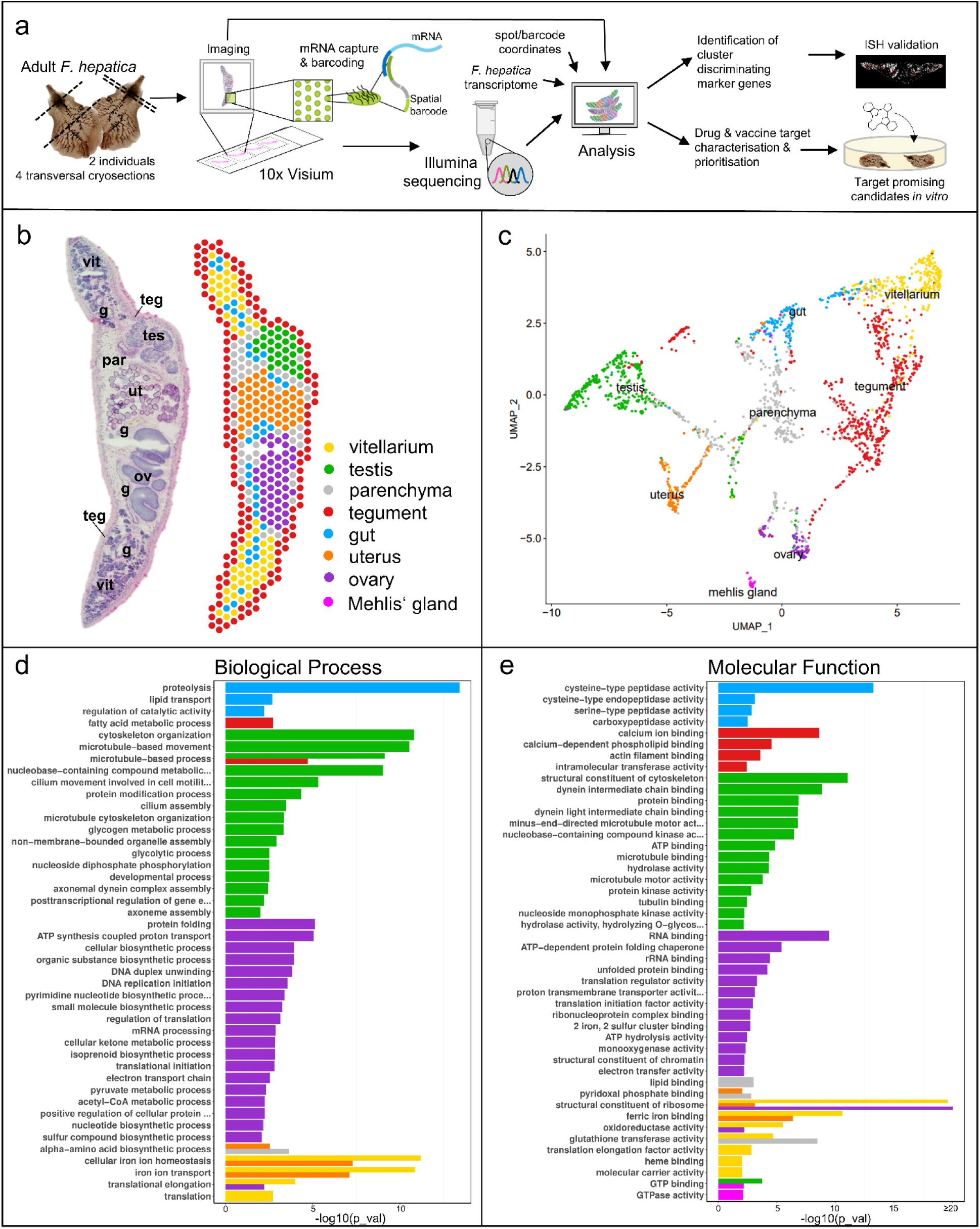
A spatial transcriptome of *F. hepatica* cross-sections. (**a**) Scheme describing the experimental workflow: Four cryosections of two adult liver flukes were placed on a 10x Visium Spatial Gene Expression slide, stained and imaged. mRNA release and barcoding were performed according to the 10x Visium protocol. During the analysis, all transcripts were mapped back to their corresponding spots on the slide and annotated using the reference transcriptome. Clustering was carried out to identify transcriptionally distinct tissues and tissue-specific markers. Selected markers were validated by *in situ* hybridization (ISH). The dataset was then used to explore spatial expression profiles of drug and vaccine candidates. One promising candidate was finally targeted with a small-molecule compound *in vitro*. (**b)** H&E stained tissue section and corresponding spatial projection of 412 mRNA-binding spots covered by this tissue section. Clusters are colored and labelled. g: gut, ov: ovary, par: parenchyma, teg: tegument, tes: testis, ut: uterus, v: vitellarium. Please note: There is no Mehlis’ gland in this tissue section. **(c)** Uniform Manifold Approximation and Projection (UMAP) of 2020 spots derived from four different *F. hepatica* cross-sections. Clusters are colored and labelled according to (b). See Figure S1 for spatial projections of sections not shown in (b). (**d,e)** Gene ontology analysis of marker genes (top 75% per cluster) revealed characteristic biological processes (d) and molecular functions (e) for each cluster. Bars for individual clusters are colored according to legend and labelling in (b) and (c). For STRING analyses of marker genes of the ovary and testis cluster see Figure S2.

All sections together covered a total of 2,020 mRNA-binding spots, each coated with millions of barcoded oligonucleotides. We captured a median of 2,192 genes and 6,138 UMIs per spot (Figure S1, Table S1). In total, over all spots, we detected transcripts of 9,847 different genes, constituting 79.3% of all gene transcripts in the *F. hepatica* genome (PRJNA179522)^15^. We then used Seurat^16,17^ to perform clustering and to identify transcriptionally distinct tissues and their marker genes. In this way, we received individual clusters representing eight tissues: tegument (561 spots), gut (154 spots), parenchyma (410 spots), vitellarium (279 spots), uterus (134 spots), ovary (92 spots), testis (354 spots) and Mehlis’ gland (36 spots) (Figures 1b, 1c, S1c-S1f). Due to the given resolution of the Visium approach (55 µm spot diameter, 100 µm spot-to-spot distance^18^), cell types that are spatially close to each other were combined in one cluster. This applies, for example, to the tegument cluster, which includes all subtegumental cells, i.e., muscle cells and some neurons in addition to tegumental cytons.

Gene Ontology (GO) enrichment analysis of marker genes revealed characteristic biological processes and molecular functions for each cluster (Figures 1d & 1e). For instance, the gut cluster exhibited a significant enrichment in genes associated with “proteolysis”. Testis and ovary clusters were enriched in genes involved in "microtubule-based movement” and biosynthesis, respectively. These analyses suggested that each cluster is molecularly distinct and that our dataset is capable to display the different biological functions of different tissue types. The tissue-specificity of selected marker genes was confirmed with *in situ* hybridization (ISH) as an independent method (Figures 2, 4, S3).

**Figure 2.**
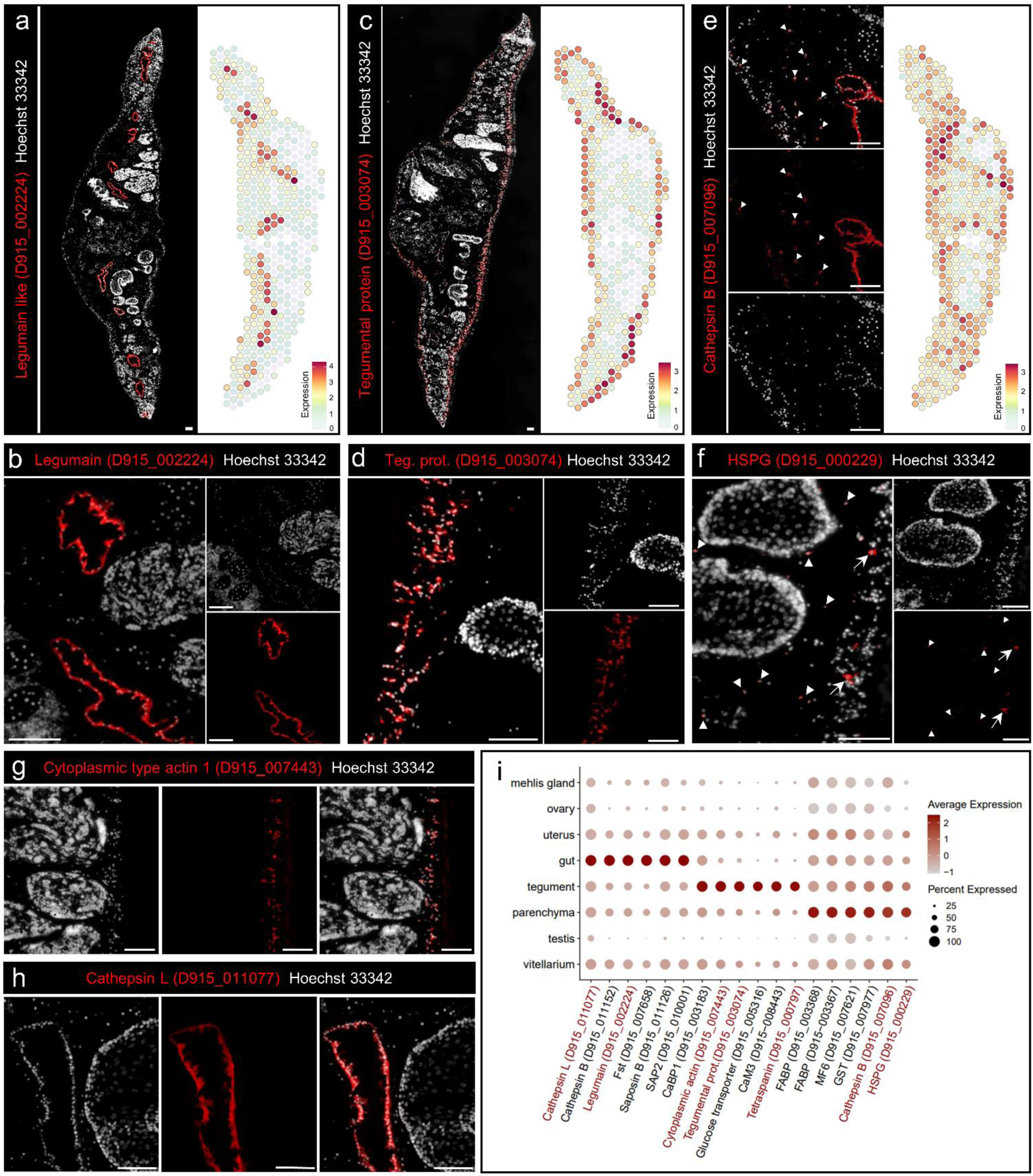
Spatial expression of selected intestinal, tegumental and parenchymal marker genes. **(a,c)** Fluorescent *in situ* hybridization (FISH) overview (left) and spatial projection (right) of the gut marker legumain (D915_002224) (**a**) and a tegumental EF hand domain containing protein (D915_003074) (**c**). (**b,d)** Magnification of (a) and (c), respectively. **(e)** FISH (left) and spatial projection (right) of cathepsin B (D915_007096). This cathepsin B is expressed in the worm parenchyma as well as in the gut, white arrowheads indicating positive parenchymal cells. (a,c,e) Expression level encoded by color (grey = low, red = high). **(f)** FISH of basement membrane-specific heparan sulfate proteoglycan core protein (HSPG) (D915_0000229). HSPG is expressed in small cells within the worms’ parenchyma (white arrowheads) as well as in larger cells below the tegument (white arrows). **(g)** FISH of cytoplasmic type actin 1 (D915_007443). **(h)** FISH of cathepsin L (D915_011077). (a-h) Scale = 100 µm. **(i)** DotPlot showing expression profiles of selected tissue markers of the gut, tegument or parenchyma cluster. Dot color encodes the average expression level (scaled) across all spots within a cluster. Dot size encodes the percentage of spots within a cluster that have captured this transcript. Please note: While spatial plots (a, c, e) are shown for only one representative section, the DotPlot includes expression data from all four tissue sections in the dataset. Genes labelled in red were validated by FISH. See Figure S3.1a for FISH of tetraspanin (D915_007977).

**Figure 3.**
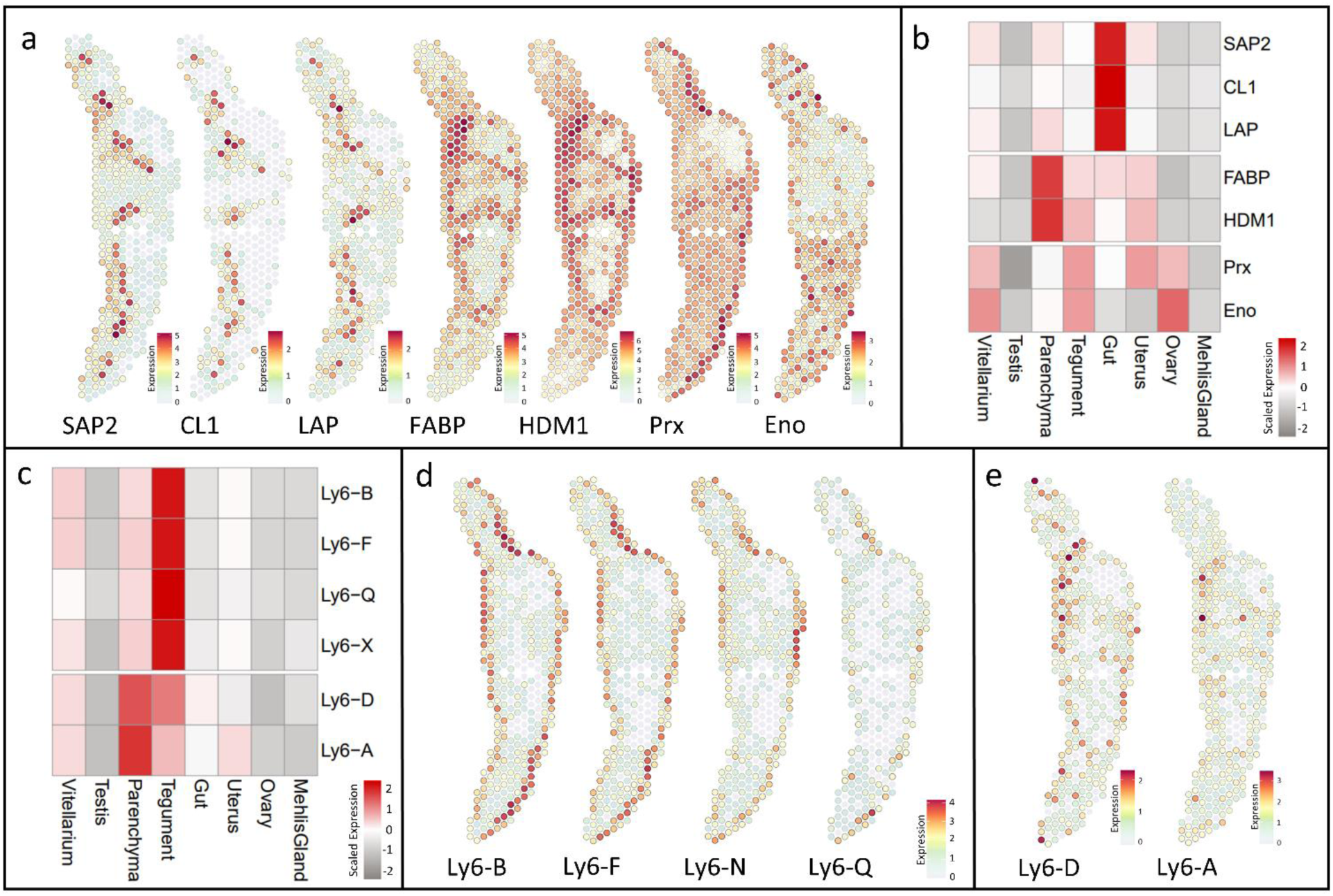
Vaccine candidates are expressed in gut, tegument and parenchyma. (**a**) Spatial projections showing expression patterns of saposin-like family protein 2 (SAP2, D915_010001), cathepsin L1 (CL1, D915_005527), leucine aminopeptidase (LAP, D915_002812), fatty acid binding protein Fh15 (FABP, D915_003367), helminth-defense molecule 1 (HDM1/ MF6, D915_007621), peroxiredoxin (Prx/TSA, D915_003729) and enolase (Eno, D915_008715). **(a,d,e)** Expression level encoded by color (grey = low, red = high). **(b,c)** Heatmap showing the average expression per cluster of genes shown in (a) and (d,e), respectively. Expression values were centered and scaled for each row (each gene) individually. Please note: While spatial plots (a, d, e) are shown for only one representative section, the heatmap includes expression data from all four tissue sections in the dataset **(d,e)** Spatial projections showing expression patterns of tegumental and parenchymal *F. hepatica* Ly6 proteins. **(d)** Tegumental Ly6 proteins: Ly6-B (D915_008996), Ly6-F (D915_008863), Ly6-N (D915_007373), Ly6-Q (D915_008394). **(e)** Ly6 proteins with prominent expression in the parenchyma: Ly6-D (D915_006706), Ly6-A (D915_001097). Figure S4a for Ly6 proteins not shown in (c)-(e).

**Figure 4.**
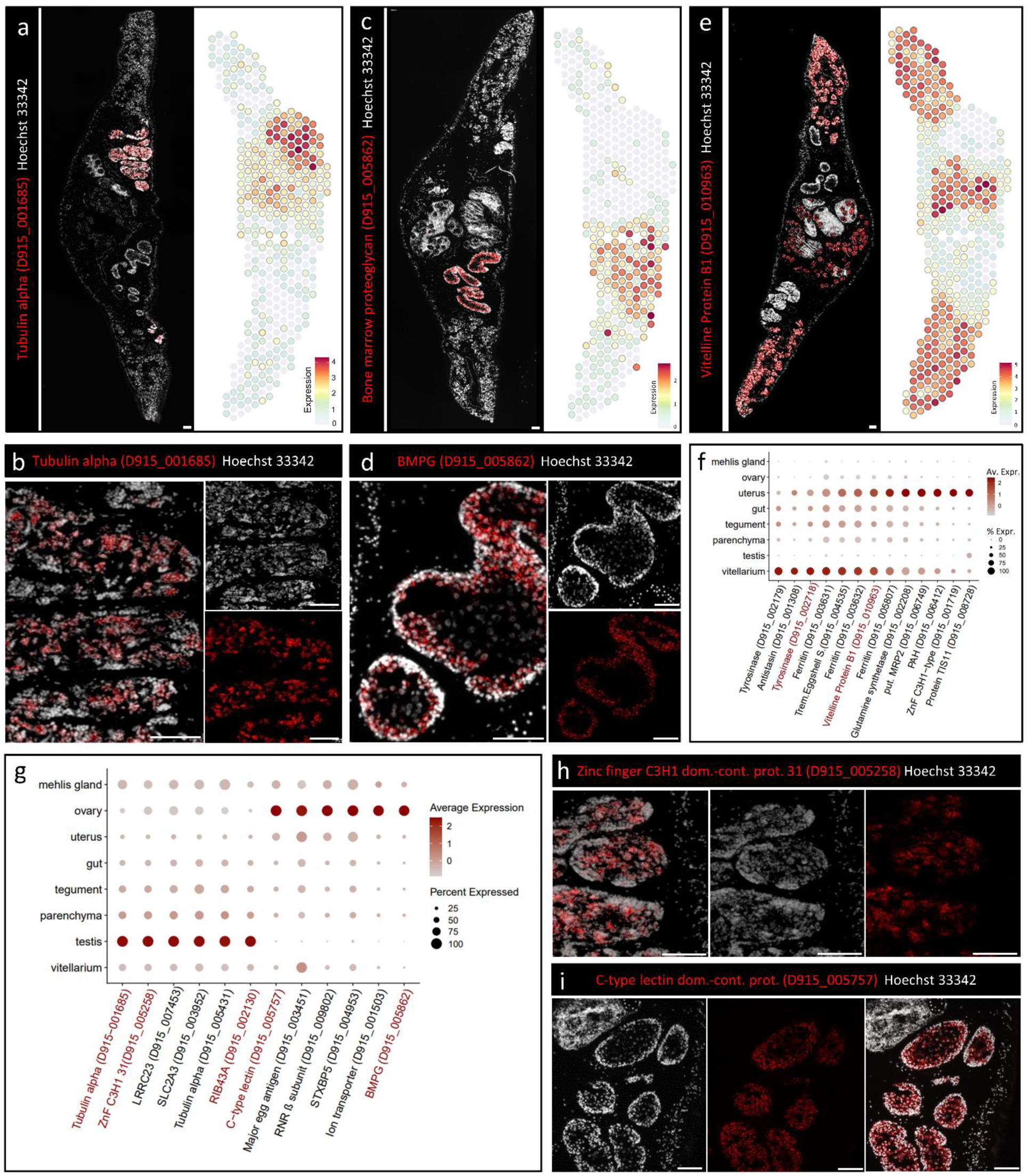
Spatial expression of marker genes for liver fluke reproductive organs. **(a,c,e)** Fluorescent *in situ* hybridization (FISH) overview (left) and spatial projection (right) of a testis specific tubulin alpha (D915_001685) (a), the ovary marker bone marrow proteoglycan (BMPG, D915_005862) (c), and the vitelline protein B1 (D915_010963) (e). Expression level encoded by color (grey = low, red = high). **(b,d)** Magnified view of FISH stainings for genes shown in (a) and (c), respectively (different tissue section, same experiment). **(f,g)** DotPlot showing expression profiles of selected marker genes for vitellarium and uterus (f) or testis and ovary (g). Dot color encodes the average expression level (scaled) across all spots within a cluster. Dot size encodes the percentage of spots within a cluster that have captured this transcript. Please note: While spatial plots (a, c, e) are shown for only one representative section, the DotPlot includes expression data from all four tissue sections in the dataset. Genes labelled in red were validated by ISH. See Figure S3.1b & S3.1c for ISH of RIB43A (D915_002130) and tyrosinase (D915_002718)**. (h)** FISH of Zinc finger C3H1 domain-containing protein 31 (D915_005258). **(i)** FISH of C-type lectin domain-containing protein (D915_005757). (a-e,h,i) Scale = 100 µm.

### Marker genes reflect functional complexity of gut, tegument and parenchyma

Trematodes have developed several functional and morphological adaptations to a parasitic lifestyle. For instance, the fluke’s gut is well equipped to digest host erythrocytes and hemoglobin and thereby provide amino acids needed for the production of 25,000 eggs per day^19^. This digestive function was well reflected by 17 digestive enzymes, mainly proteases and hydrolases, among the top 50 marker genes (Table S2). Expression of the two cysteine proteases legumain (D915_002224) and cathepsin L (D915_011077) could be specifically allocated to the intestinal epithelium via ISH (Figures 2a, 2b & 2h).

The tegument is another remarkable feature of parasitic flatworms. Similar to the gut, it serves absorptive functions, but it also acts as protective layer at the host-parasite interface. Related to nutrient import, expression of a glucose transporter (D915_005316) and an amino acid transporter (D915_001928) was enriched within the tegument cluster. Furthermore, analysis of annotated GO terms for this cluster showed an enrichment of molecular functions associated with “calcium ion binding” (Figure 1e). This GO term is represented by three annexins, one calmodulin 3, one alpha-actinin and seven EF hand domain containing proteins (Table S2). ISH detected transcripts of the EF hand domain-containing protein D915_003074 within groups of cells sitting below the body wall musculature with cytoplasmic protrusions towards the syncytial layer (Figure 2d). A similar expression was found for cytoplasmic type actin (D915_007443) and a tetraspanin family protein (D915_000797) (Figures 2g & S3.1a), two important proteins related to cytoskeleton and membrane structure.

Furthermore, our dataset allowed to shed light on the gene expression characteristics of parenchyma, a specialized tissue in flatworms embedding all other organs. It functions both as a flexible cytoskeleton and as a site for synthesis, storage, and transport of nutrients^20,21^. GO term analysis suggested lipid, amino acid and drug metabolism (“glutathione transferase activity”) as main functions (Figures 1d & 1e). Top markers such as fatty acid-binding proteins (D915_003368, D915_003367), HDM-1 (“MF6”/D915_007621) and glutathione S-transferases (GSTs) will be further addressed below. Two newly identified markers of the liver fluke parenchyma were a heparan sulfate proteoglycan (HSPG, an extracellular matrix protein) (D915_000229) and a parenchymal cathepsin B (D915_007096) (Figure 2i, Table S2). Liver fluke HSPG possesses similarities to human HSPG2, which is a functionally diverse protein whose different domains are able to bind other matrix components, cells, LDL and growth factors ^22,23^. ISH confirmed the parenchymal expression of both HSPG and cathepsin B (Figures 2e & 2f).

Taken together, this dataset enables us to make inferences regarding the various biological functions of the liver flukes’ three largest somatic tissues, which includes the identification of new marker genes.

### Spatial expression of vaccine candidates

The liver fluke gut, tegument and parenchyma share as common feature that they are all in close interaction with the hosts’ immune system, either by direct contact or by synthesis and release of excretory/secretory (ES) products and extracellular vesicles (EVs)^12,21^. Therefore, it is no surprise that we detected several published vaccine candidates within these tissues (Figures 3a & 3b). For the intestine, these were cathepsin L1 (D915_005527)^24^, leucine aminopeptidase (“LAP, D915_002812)^25^ and the saposin-like family protein 2 (SAP2, D915_010001)^26^. The helminth-defense molecule 1 (HDM1/MF6, D915_007621)^27^ and the fatty acid binding protein Fh15 (FAPB, D915_003367)^28^ both depicted strongest expression in the worm’s parenchyma. Peroxiredoxin/ thioredoxin peroxidase (Prx/TSA, D915_003729)^29^ as well as enolase (Eno, D915_008715)^30^ displayed a broad expression across multiple tissues, but clearly included the tegument next to reproductive tissues.

Next, we focused on CD59-like proteins of the Ly6 family, which are thought to be involved in parasite-host interactions and have recently been proposed as vaccine candidates for *Fasciola spp*.^31^. We first performed WormBase ParaSite BLASTp to identify orthologues of *F. gigantica* Ly6 proteins within the *F. hepatica* genome (PRJNA179522). In total, we found 19 FhLy6 proteins (see Table S4 for details). We then searched our spatial transcriptome for all family members. Thereby, we identified four tegumental (Ly6-B (D915_008996), Ly6-F (D915_008863), Ly6-N (D915_007373), Ly6-Q (D915_008394)) and two mainly parenchymal Ly6 proteins (Ly6-A (D915_001097), Ly6-D (D915_006706)) (Figures 3c-3e). Thus, spatial transcriptomics could extend our knowledge on expression of different vaccine candidates and highlighted the role of intestine, tegument and parenchyma in parasite-host interaction.

### Male and female gonads: A highly dynamic germ cell production machinery

Another remarkable adaptation of trematodes to the parasitic lifestyle is an exceedingly high fecundity, which allows the parasites to spread efficiently among hosts. Liver flukes are hermaphrodites and male and female reproductive organs make up a large proportion of the adult fluke. The high metabolic activity and rapid rate of cellular differentiation and turnover in these organs was well reflected in the high number of UMIs and genes captured on corresponding spots. The testis and ovary cluster showed the highest transcriptional activity and the largest number of expressed genes compared to all other tissues (Figures S1c-S1h).

GO-term and STRING analysis for the ovary showed that those transcripts correspond to proteins involved in a variety of biosynthetic processes, in particular (ribo-)nucleotide and small molecule synthesis, DNA replication and translation (Figures 1d & S2c). The development of early mammalian embryos depends on the biosynthetic activity and stored transcripts of the oocyte^32^. Our data suggest this is also true for oocytes of *F. hepatica*.In addition, we identified two C-type lectins of unknown function with distinct spatiotemporal expression during oocyte differentiation. The liver fluke ovary is structured in a way that oogonia and early primary oocytes reside in the periphery of the ovarian tubule while late primary oocytes are found in the center^33^. By ISH, we showed that the bone marrow proteoglycan (BMPG, D915_005862) was predominantly expressed in early primary oocytes, but far less in late primary oocytes (Figure 4d), while D915_005757 transcripts were also detected in late primary oocytes (Figure 4i).

The production machinery of the testis is clearly directed towards one goal: the production of large numbers of motile spermatozoa. STRING analysis for marker genes of the testis cluster displayed a tight network of protein-protein associations (Figure S2a). The network was functionally enriched in numerous terms associated with “Cytoskeleton”, “Microtubule”, “Cilium” and “Axoneme”. Markers included several alpha and beta tubulins and the flagellar ribbon protein RIB43A (D915_002130) (Figure 4g, Table S2). ISH demonstrated that the alpha tubulin D915_001685 and RIB43A were expressed in almost all stages of spermatogenesis, apart from spermatogonia, which are located in the periphery of the testicular tubules^33^ (Figures 4b & S3.1b).

To ensure successful germ cell formation, cellular processes must be tightly regulated. Against this background, we noticed enriched expression of seven and eight different ovarian and testicular Zinc finger proteins, respectively (Table S2). These included several C2H2 class proteins and CCCH domain-containing proteins (D915_005258, D915_003685), which stand out among zinc fingers as they bind RNA not DNA and thereby regulate RNA metabolism^34^. Indeed, the CCCH zinc finger D915_003685 was part of a STRING subnetwork of ovary marker genes functionally enriched in genes associated with “mRNA processing” and “RNA splicing” (Figure S2d). Additionally, ISH was used to confirm the expression of the Zinc finger CCCH domain-containing protein 31 (D915_005258) in the testis (Figure 4h).

### The parasite’s egg production apparatus: from vitellarium to uterus

Trematode eggs are composite eggs, made up of an oocyte and about 30 vitelline cells surrounded by an egg shell^35^. The lateral margins of adult liver flukes are filled with vitelline follicles, which provide vitelline cells that synthesize proteins and enzymes needed for eggshell formation^36^. This includes eggshell proteins, tyrosinases and multiple iron-binding proteins such as ferritins and myoglobin. Spatial transcriptomics not only confirmed vitellarial expression, but could also shed light on the type of vitelline cell stages that express particular genes. While the tyrosinase 1 and 2 (D915_002197, D915_002718) were expressed exclusively within vitelline cells in the vitellarium (stages S1-S3), other markers, such as the vitelline protein B1 were also expressed in mature vitelline cells (stage S4) found within eggs in the uterus (Figures 4e, 4f & S3.1c).

The fluke’s uterus is typically filled with eggs in different stages of eggshell formation, which was also the case for the specimens of our spatial transcriptomic analysis. Since each egg consists of dozens of vitellocytes^35^, the uterus cluster shares several marker genes with the vitellarium cluster (Figure 4f). Nevertheless, we were able to identify some genes that are expressed almost exclusively within the uterus and might be involved in early embryogenesis. This includes the glutamine synthetase D915_002208, the phenylalanine 4-monooxygenase D915_006412 and the mRNA-binding protein TIS11 (D915_008728) (Figure 4f).

The Mehlis’ gland or the so-called shell gland is another part of the female reproductive apparatus and a unique feature of the reproductive system of parasitic flatworms^37^. We achieved manual subclustering of this organ into a dorso-lateral and ventral region, which express distinct marker genes associated with two gland cell-types (Figure S3.2, Table S3) only described morphologically so far^33^.

Taken together, our spatial atlas provided unprecedented insight into gene expression of the diverse organs characteristic for the unique trematode reproductive system.

### Spatial transcriptomics suggests functional subsets of the *F. hepatica* β-tubulin family

Beta-tubulins are of research interest as they are molecular targets of triclabendazole, a benzimidazole and the drug of choice to treat fasciolosis^11^. All six β-tubulins (isoform 1-6) of *F. hepatica* previously described in literature^38^ were found in the spatial transcriptome.

Additionally, we identified two new, complete β-tubulin isoforms (β-tub7 (D915_000542) and β-tub8 (D915_000946)) in the *F. hepatica* genome (PRJNA179522) in WormBase ParaSite (see Figures S5a-S5c for details). β-tubulin 7 was not present in the spatial transcriptomic dataset. For the remaining seven isoforms, our data revealed characteristic expression patterns (Figures 5a-5c). Among the already known β-tubulin isoforms, we were able to identify two major subsets based on their spatial expression. β-tubulin 1, 5, and 6 were almost exclusively expressed in the testis (Figure 5a), while isoforms 2, 3a, and 4 were expressed in a broader range of tissues (Figure 5b). Isoforms 3a and 4, as well as the newly discovered β-tubulin isoform 8, showed a peak expression in the flukes’ ovary (Figure 5b). β-tubulin 2 in contrast was predominantly expressed in non-reproductive tissues, with highest expression in the tegument (Figure 5b). The expression level of β-tubulin 3b was relatively weak compared with the other isoforms and could not be confidently assigned to specific tissues (Figure S4e). A similar grouping into testis-associated isoforms and isoforms expressed elsewhere was also observed for α-tubulin isoforms 1-5 (Figures S4b – S4d).

**Figure 5.**
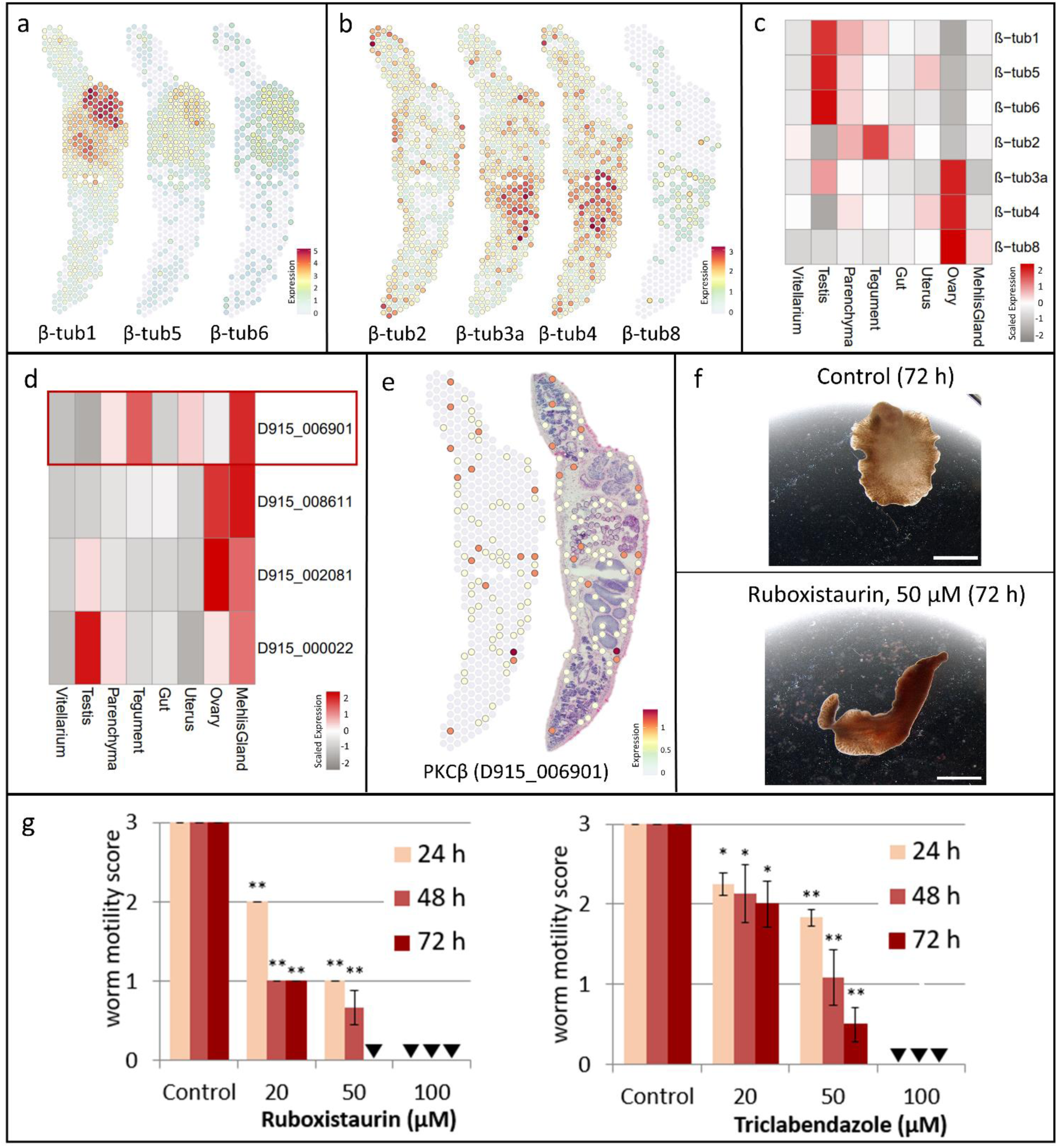
Spatial transcriptomics allows drug target characterization and prioritization. **(a,b)** Spatial projections showing expression patterns of *F. hepatica* β-tubulins. **(a)** β-tubulin isoforms with predominant expression in the testis: β-tub1 (D915_007398), β-tub5 (D915_003963), β-tub6 (D915_008457) **(b)** β-tub2 (“D915_002311), β-tub3a (D915_004911), β-tub4 (D915_001342), β-tub8 (D915_000946). For spatial projections of β3b and alpha-tubulins see Figure S4b, S4c & S4e. **(c)** Heatmap showing the average β-tubulin isoform expression per cluster. Expression values were centered and scaled for each row (each gene) individually. **(d)** Heatmap showing the average expression of different protein kinase C (PKC) genes per cluster. Expression values were centered and scaled for each row (each gene) individually. Red rectangle marks PKCβ with tegumental expression. **(e)** Spatial projection only (left) and overlay with H&E stained tissue section (right) showing expression of PKCβ (D915_006901). Several positive spots can be seen along the tegument of the parasite. For spatial projections of other PKCs see Figure S4f. (a,b,e) Expression level encoded by color (grey = low, red = high). Please note: While spatial plots (a, b, e) are shown for only one representative section, heatmaps include expression data from all four tissue sections in the dataset. **(f,g)** Adult flukes were treated for 72 h with different concentrations of the PCKβ isoform-specific inhibitor ruboxistaurin (20-100 µM) or triclabendazole as positive control. Motility was assessed every 24 h. Control worms were treated with the inhibitor solvent, DMSO. Representative images are shown in (f) and motility scores for all time-points and concentrations in (g) (score 3 = normal, 2= reduced, 1= severely reduced, 0 = no motility). See also Video S1 & S2. Data represent the mean±SEM of 2-3 independent experiments (4-6 flukes per group). Significant differences to controls are indicated with *p<0.05 **p<0.01 (Mann-Whitney test). Triangle indicates a score of zero. Scale bars correspond to 5 mm.

### Exploring the spatial expression of PKCs to assess their drug target potential

With the rationale that new drug targets may be particularly found among tegumental or intestinal proteins (organs vital for the parasite), we sought to survey the tegument cluster for putative targets. Based on our previous research on protein kinases of *F. hepatica*^13^, we focused on this particular class of druggable proteins and more specifically, on family members of protein kinase C (PKC), cell signaling enzymes with known importance for helminth physiology^39,40^. Among four PKC genes that we identified in the genome of *F. hepatica,* one, a PKCβ (D915_006901), showed notable expression in the tegument cluster (Figure 5d & 5e). Based on an amino acid identity of 73.36% between the catalytic domains of *F. hepatica* and human PKCβ (Figure S5e), we made use of the highly isoform-specific human PKCβ inhibitor ruboxistaurin (LY333531)^41^ to test whether targeting of PKCβ causes anthelminthic effects. Indeed, treatment with 50 µM ruboxistaurin killed adult *F. hepatica* within 72 h of *in vitro* culture, a potency comparable to the gold standard triclabendazole (Figure 5f & 5g, Video S1 & S2). This is one clear example how the spatial transcriptome may help drug search and target prioritization.

### Spatial distribution of GSTs supports a specialized role of the parenchyma in detoxification

Glutathione S-transferases (GSTs) are among the molecules that defend the parasite against immune-induced damage and may also mediate cellular detoxification of drugs^42^. For *F. hepatica*, GSTs out of four classes (Mu, Sigma, Omega and Zeta) have been identified and characterized by biochemistry and bioinformatics^43–45^. Our work complements these findings with information on the spatial expression of 11 cytosolic (6x Mu, 2x Sigma, 2x Omega, 1x Zeta) and two microsomal GSTs. Phylogenetic analysis of their sequenes (retrieved from the “St. Louis” *F. hepatica* genome, PRJNA179522) together with human and known *Fasciola* GST sequences confirmed isoform assignment (Figure S6a).

For the large group of Mu class GSTs, spatial transcriptomics revealed a common characteristic feature: a preference for parenchymal expression (Figure 6a, 6e & S4g). One exception was GST-Mu5 (D915_002901)^45^, which was mainly expressed within the tegument (Figures 6a & 6e). The two Sigma class GSTs present in the dataset clearly diverged in their spatial expression patterns. GST-S1 (D915_001201) expression in the vitellarium clearly exceeded that in all other organs, while expression of GST-S2 (D915_006844) was highest in the parenchyma (Figures 6b & 6e). Diverging spatial expression patterns were also obvious for the two omega-class GSTs. While GST-O1 (D915_001421) was mainly expressed in the vitellarium and uterus and somewhat less in the ovary, GST-O2 (D915-001777) was predominantly expressed in the parasite’s tegument (Figures 6c & 6e). These results support a role for GST-O1 within the reproductive system, but suggest a different role for GST-O2 in protecting the parasite’s barrier to the host. Finally, we found two microsomal GSTs with expression patterns complementing each other. While GST-m1 (D915_002950) was expressed in almost all tissues except testis, GST-m2 (D915_007840) showed strong expression within spots assigned to the testis cluster (Figures 6d & 6e). In conclusion, most organs are characterized by a specific set of GSTs, which may reflect different needs of the tissues with respect to detoxification. Particularly noteworthy is the parenchyma, which expresses the greatest diversity of GSTs (Figure 6e).

**Figure 6.**
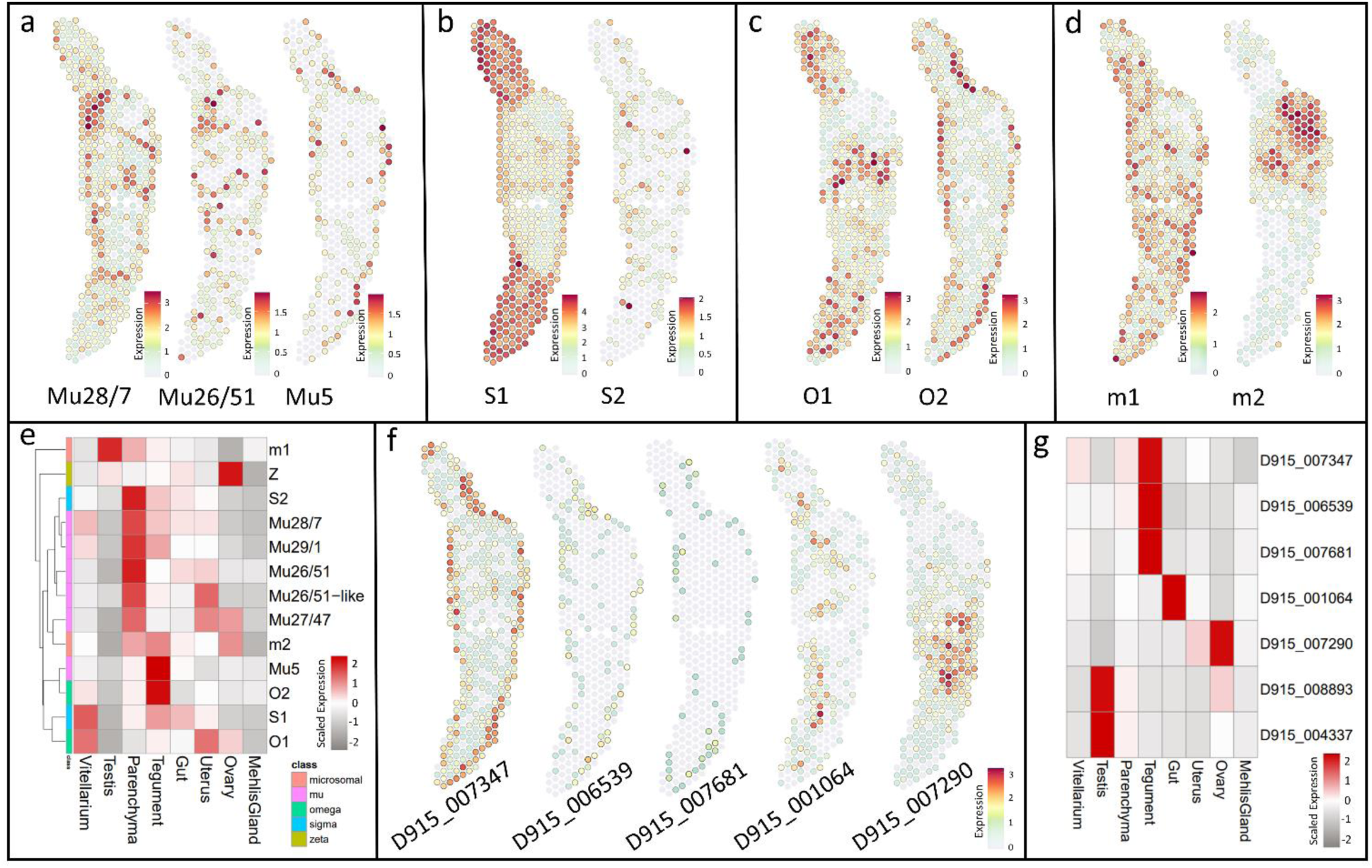
Tissue specific expression of detoxifying enzymes (glutathione S-transferases) and efflux transporters (ABC transporters) **(a-b)** Spatial projections showing expression patterns of *F. hepatica* glutathione S-transferases (GSTs). (**a)** Mu-class GSTs: Mu28/7 (D915_008526), Mu26/51 (D915_008966), Mu5 (D915_002901) **(b)** Sigma-class GSTs: S1 (D915_001201), S2 (D915_006844) **c** Omega-class GSTs: O1 (D915_001421), O2 (D915_001777) **(d)** microsomal GSTs: m1 (D915_007840), m2 (D915_002950). **(e)** Heatmap of GSTs showing the average expression per cluster of genes. Expression values were centered and scaled for each row (each gene) individually. Rows are clustered. Annotation column indicates GST class: mu (pink), microsomal (orange), sigma (blue), omega (green), zeta (yellow). Also includes additional GSTs not shown in (a-d): Mu27/47 (D915_010266), Mu29/1 (D915_007977), Mu26/51-like (D915_008366), Zeta (D915_006391), see Figure S4g & S4h for spatial projections. **(f)** Spatial projection showing expression patterns of selected *F. hepatica* ABC-B transporters. **(g)** Heatmap of ABC-B transporters showing the average expression per cluster of genes shown in (f) as well as two testis-associated ABC-B transporters. Expression values were centered and scaled for each row (each gene) individually. See Figure S4i for spatial projections of ABC-B genes not shown in (f). (a-d,f) Expression level encoded by color (grey = low, red = high). Please note: While spatial plots (a-d, f) are shown for only one representative section, heatmaps include expression data from all four tissue sections in the dataset.

### Tissue-specific expression of *F. hepatica* ABC-B transporters

ABC transporters represent another interesting protein family of which some members help the parasite to defend against toxic products, most likely including drugs such as triclabendazole^11^. By phylogenetic modelling of 27 *F. hepatica* ABC transporter sequences together with human and *C. elegans* ABC transporters from all subfamilies (A,B,C,D,E,F,G), we verified in total: 4 ABC-A, 12 ABC-B, 3 ABC-C, 2 ABC-D, 1 ABC-E, 2 ABC-F and 3 ABC-G members (Figure S6b).

As subfamily B is particularly interesting with regard to drug resistance^11^, we have focused on this subfamily in the further course. For seven family members we identified an association to specific organs, while expression of the others was rather weak and dispersed (Figures 6f, 6g & S4i). Most striking was the expression of four genes (D915_007347 D915_006539, D915_007681 and D915_001064) in tissues at the host-parasite interface (Figures 6f & 6g). Particular noteworthy is the strong expression of D915_007347 in the tegument (Figures 6f & 6g). D915_001064, on the other hand, was the only ABC-B transporter that was mainly expressed in the intestine. Expression of other ABC-B genes, above all D915_007290, was localized to the worm’s gonads. While D915_007290 was strongly expressed in the ovary, the expression of D915_008893 and D915_04337 was rather weak and mainly associated with the testis of the worm (Figures 6g & S4i). Thus, as for GSTs, we could reveal a tissue-specific expression of selected ABC transporters, pointing to distinct biological roles.

## Discussion

The biology of parasitic metazoans, including the function of their various tissues, is still poorly understood. Their sheer size and cell number, complex life cycles, and a lack of molecular tools are turning parasite research into an experimental challenge. Nevertheless, the need for novel therapies against these pathogens has steadily driven research in this area. Now, we are able to provide a comprehensive 2D expression atlas of a multicellular parasite, the common liver fluke *F. hepatica*.

### Technical advancement for spatial expression analysis of parasite genes

Various approaches have been used in the past to analyze the proteome or transcriptome of body regions, individual organs and tissues to gain molecular insights into the complex biology of these organism. These methods, however, usually focused on a small number of selected organs. For example, the *F. hepatica* and schistosome tegument proteome were studied using different physical and enzymatic preparation techniques in order to detach the tegumental syncytium from the worm surface^46,47^. Hahnel et al.^48^ developed a digestion protocol to isolate reproductive organs of male and female schistosomes, which were then further analyzed using RNAseq. Furthermore, an RNA tomography approach combined with laser capture microdissection (LCM) enabled insights into the spatial and tissue transcriptome of the *Brugia malayi* head region^49^. LCM technology has also been used to excise and analyze the liver fluke gut and tegument proteome^50^ as well as the transcriptome of *Schistosoma japonicum* intestine, vitellarium and ovary^51^.

The increasing number of parasite genomes and the development of ground-breaking “omics” technologies in recent years have also opened up new opportunities for parasite research^12,52^. The examples of *Schistosoma mansoni* and *B. malayi* demonstrate that single-cell transcriptomics allows us to identify and characterize the cellular composition of parasites^53–55^. However, what scRNAseq cannot do is to show the spatial arrangement of the cells’ gene expression in the parasite. To do so, classical *in situ* hybridizations or immunohistochemistry are still necessary, which are associated with the time-consuming production and testing of transcript- or protein-specific riboprobes or antisera. In contrast, a spatial transcriptomics dataset, as presented here, allows to assess the whole spatial transcriptome of all tissues within a tissue section at once. And it does so without the need for time-consuming preparation steps, as in the methods described above, and without being forced to select individual organs of interest in advance. Even though a spatial transcriptome based on mRNA-capturing spots does not (yet) have the same cellular resolution as a scRNAseq dataset or an *in situ* hybridization, it allows a quick and easy assessment of the whole spatial transcriptome of a tissue section. *In situ* hybridizations can thus be partially replaced, especially when an overview of the spatial gene expression of many different genes or even a whole gene family is required. Only if there is interest in a more precise or even (sub)cellular localization of individual transcripts, a method with higher resolution is still necessary.

Our tissue atlas provides a comprehensive spatial gene expression map of adult *F. hepatica*. It allowed the characterization of eight distinct clusters, corresponding to distinct tissues. In case of Mehlis’ gland and vitelline cells we were even able to characterize specific cell types and stages, respectively. We used classical ISH experiments to validate marker genes for each of the tissues. In doing so, we confirmed known tissues markers such as cathepsin L within the gut^56^ and calcium binding proteins within the tegument^57,46^, but also uncovered new molecular markers for several tissues (e.g. HSPG in the parenchyma, a testis-specific C3H zinc finger). This proofs that our data reflect the true localization of transcripts.

For the most studied trematode, *S. mansoni*, spatial transcriptomics data are not available, but we can make comparisons to existing scRNAseq data to reveal tissue-specific gene signatures that may be evolutionary conserved among parasitic flatworms. For instance, among the marker genes highlighted in our study, parenchymal cathepsin B, ovarian bmpg and vitelline tyrosinase 1 and 2 have also been found as tissue-specific marker genes in *S. mansoni*^54,53^.

### Utilizing spatial gene expression data for applied research

To test the power of our spatial atlas, we focused on genes of high interest for the field of parasitology: vaccine candidates and four gene families associated with drug action or resistance. Tegumental and intestinal proteins appear as particularly attractive vaccine and drug targets thanks to their exposure towards the host^12^. Thus, the spatial transcriptome, covering thousands of genes expressed within these tissues, can help prioritizing candidate proteins in the future. As proof-of-concept, we could identify a tegumentally expressed PKCβ and potent anthelminthic activity of a PKCβ inhibitor, ruboxistaurin. This compound was used in several clinical studies addressing diabetic retinopathy, was safe in patients and orally bioavailable^58^. This motivates follow-up studies on ruboxistaurin as drug candidate against *F. hepatica* infection. Furthermore, we were able to visualize the spatial expression of 16 Ly6 proteins, seven β-tubulins, 13 glutathione S-transferases and 12 ABC-B transporters.

*In silico* characterization of the Ly6 protein family has recently revealed several new members and a high diversity of this gene family in *F. hepatica* ^31^. In humans, the Ly6 protein CD59 can prevent formation of the complement membrane attack complex at the cell surface^59^. Whether a similar defense against immune attack may hold true for liver flukes is still unknown^31^. However, with reference to promising preliminary vaccination trials using *S. mansoni* Ly6 proteins^60^, this gene family has recently been suggested as a vaccine candidate for *Fasciola spp*.^31^. To be able to prioritize and select certain Ly6 proteins for vaccine trials, knowledge on the spatial expression of the different family members would be highly beneficial. Until now, only Ly6-Q (D915_008394) could be detected in the tegument proteome of adult parasites^46^. We were now able to extend the list of potential vaccine candidates to at least four tegumental (Ly6-B, Ly6-F, Ly6-N, Ly6-Q) and two parenchymal Ly6 proteins (Ly6-A, Ly6-D). Three others (Ly6-M, Ly6-O, Ly6-R) were detected predominantly in the testis, an organ not vital for the parasite, and therefore appear not to be the best vaccine candidates. Interestingly, *S. mansoni* homologues of FhLy6-Q and FhLy6-B were also localized in the tegument^61,62^, suggesting a conserved but unknown function of these proteins in the tegument of different trematode species.

Differential expression of tubulin isoforms among different cell types is a common phenomenon in eukaryotes, suggesting that some isoforms are better suited for certain cell type-specific functions^63^. The β-tubulin isoforms of *Fasciola spp.* can be divided into two functional subsets based on their basal expression in different life stages^64^. Three β-tubulin isoforms (β2, β3, β4) showed high constitutive expression, implying a “housekeeping” role for general microtubule structure and function, whereas the remaining isoforms (β1, β5, β6) displayed a marked upregulation in immature and adult flukes, indicating a more specialized, but so far undetermined role^64,12^. Our spatial transcriptome now revealed that tubulin isoforms in *F. hepatica* also display distinct spatial expression patterns. Isoforms with constitutive expression at the life stage also showed broader expression in various tissues, including the parenchyma, the tegument and the ovary of the fluke, while those isoforms that were upregulated in immature and adult flukes were all found to be expressed within the flukes’ testis. This implicates a specialized role of those isoforms in spermatogenesis. The expression of β-tubulin 7 was very low in adult parasites^64^, which fits with the fact that we didn’t detect it in our spatial transcriptome. Finally, the newly identified β-tubulin 8, which we found to be predominantly expressed in the ovary of adult flukes, seems to be also relevant (i.e. highly expressed) in other life stages, namely eggs, metacercariae and NEJs^64^, suggesting multiple functions during the parasite’s life cycle.

To date, the interaction of TCBZ and other benzimidazole compounds with liver fluke β-tubulins has not been fully understood^11^. Also which of the different β-tubulin isoforms are targeted by TCBZ remains unclear. It has been shown that some isoforms are more likely to bind benzimidazole compounds, while others are more refractory^65^. Merging existing knowledge on the histological changes in testis, ovary and tegument of TCBZ-treated *F. hepatica*^33,66^, with the new spatial expression data leads us to the conclusion that TCBZ may attack numerous β-tubulin isoforms, with β-tubulin 2 and β-tubulin 3 as most likely candidates implicated in tegumental damage.

### Spatial expression of ABC transporters and their potential role in drug resistance

The ABC transporter family is a large protein family with at least 27 members in *F. hepatica*. In addition to a possible role in TCBZ resistance^11^, ABC transporters may be interesting targets for developing new treatments that enhance the efficacy of existing drugs or that interfere with the physiology of the parasite^67^. However, to date, there has been almost no information on the spatial expression of ABC transporters in this parasite that would provide first insight into their biological function. We found three members of the ABC-B subfamily to be predominantly present in the fluke tegument (D915_007347, D915_006539, and D915_007681). In addition, D915_001064 was found to be mainly expressed in the intestine. Interestingly, all four have in common that they were localized in the membrane of extracellular vesicles (EV) in previous studies^50,68^. It has therefore been suggested that ABC-B transporters play a role in EV packing^68^. Our data now implies that different ABC-B transporter isoforms are involved in EV formation depending on the organ of origin.

Several ABC-B genes have been discussed in terms of their potential role in drug resistance in*F. hepatica*. One (marker-scaffold10x_157_pilon-snap-gene-0.179 = D915_004337) was located within the TCBZ resistance locus identified by Beesley et al.^69^. The other (BN1106_s3396B000087 = D915_008838) was found upregulated in a RNAseq study of a TCBZ and albendazole resistant parasite strain^70^. However, in our data both genes were primarily associated with the testis, suggesting a role in gametogenesis (e.g. protecting the germline against xenobiotics), rather than vital functions. Since we and others demonstrated that TCBZ is taken up via the tegument^71,72^, it would be reasonable to assume that mutation or increased expression of tegumental ABC transporters would favor TCBZ resistance. In our data, D915_007347 was found to be highly expressed in the tegument. A single nucleotide polymorphism (SNP) has been described for the same gene by Wilkinson et al.^73^ for a small number of TCBZ resistant *F. hepatica*, but could not be further confirmed as a resistance marker in following studies^11^. Therefore, it might be worthwhile to explore increased expression of tegumental D915_007347 in TCBZ resistant strains as an alternative mode of resistance.

### Conclusions

The aforementioned examples illustrate the utility of the dataset in exploring the spatial expression of a substantial number of genes. Thus, this dataset holds the promise of being a valuable resource for both fundamental research and drug development against the common liver fluke. Insights into the spatial expression of genes aid in deciphering their function and thus contributing to a better understanding of biology in general, and parasitism in particular. We envision these data to serve as a role-model for the investigation of other understudied and experimentally challenging multicellular parasites, improving our understanding of their complex biology and facilitating the discovery of novel therapies for these pathogens.

## Materials and Methods

### Ethical statement

Male Wistar rats RjHan:WI (*Rattus norvegicus)* (Janvier, France) were used as model hosts for *F. hepatica* infection. Rats were orally infected with 25 metacercariae at an age of 4-6 weeks and sacrifized 12-14 weeks later. Animal experiments were performed in accordance with Directive 2010/63/EU on the protection of animals used for scientific purposes and the German Animal Welfare Act. The experiments were approved by the Regional Council (Regierungspraesidium) Giessen (V54-19c20 15 h 02 GI 18/10 Nr. A16/2018).

### Preparation of parasites

All experiments were performed with an Italian parasite strain of *F. hepatica*. Metacercariae were purchased from Ridgeway Research (UK). Adult flukes were collected from the common bile duct of experimentally infected rats at 12-14 weeks p.i. Worms were kept in RPMI 1640 (Gibco, Thermo Fisher Scientific, Germany) supplemented with 5% chicken serum (Gibco, Thermo Fisher Scientific) and 1% ABAM solution (c.c.pro, Germany) at 37 °C and 5% CO2 for at least 1 h to allow clearance of gut contents. Parasites were then used for inhibitor treatments *in vitro* or embedded in O.C.T. compound (Tissue-Tek, Sakura Finetek, Germany), frozen on dry ice and stored at −80 °C until use for Visium or *in situ* hybridizations.

### Cryosectioning and section placement

We performed spatial transcriptomics using the Visium Spatial Gene Expression Solution (10x Genomics, US), which makes use of probe-coated glass slides to capture mRNA from tissue sections. Adult *F. hepatica* embedded in O.C.T. compound were used to prepare transversal cryosections of 10 μm thickness with a cryostat Microm HM525 or Epredia CryoStar NX50 (Thermo Fisher Scientific). In order to identify the tissue region of interest when creating sections, we used a quick staining (Haema Quick Stain (Diff Quick), LT-SYS, Labor + Technik Eberhard Lehmann GmbH, Germany) to stain sections every 50−100 µm and checked them with brightfield microscopy. When a region of interest was reached, the consecutive section was transferred onto the Visium Spatial Gene Expression Slide. We finally used four cryosections from two different individuals (individual 1: capture area A & B, individual 2: capture area C & D). Sections were selected to cover ovary, testis and uterus at least in one of the two sections per individual. All four sections contained tegument, vitellarium, and gut tissue. Mehlis’ gland was only present in section D. Slides with cryosections were transported on dry ice and stored in a sealed bag with desiccant at −80 °C until use.

### 10x Visium tissue processing

We used the Visium Spatial Gene Expression Slide and Reagent Kit (10x Genomics) following the manufacturer’s instructions with minor changes (for reagents see Table S5). Tissue sections were fixed in methanol, H&E-stained and imaged. Tile scanning was performed using a Leica DMi8 microscope (Leica Microsystems, Germany), equipped with a DMC2900 color camera (Leica Microsystems) and a HC PL APO 20x/0.80 objective (Leica Microsystems). Subsequently, the tissue was permeabilized to release mRNA from the tissue. Permeabilization time was previously optimized using the Visium Spatial Tissue Optimization Slide and Reagent Kit following manufacturer instructions (10x Genomics). The optimal permeabilization time was determined to be 12 minutes. Tissue permeabilization was followed by a reverse transcription, second strand synthesis and denaturation. The cDNA from each capture area was then transferred to a corresponding PCR tube to allow amplification and library construction.

### 10x Visium library preparation and sequencing

We performed qRT-PCR to determine the optimal number of PCR cycles for cDNA amplification to generate a sufficient amount of cDNA for library construction (cycler: Rotor-Gene-Q, Qiagen, Germany, reaction volume: 20 µl, protocol modified: initial denaturation step at 98 °C for 3 min; 40 cycles of: 98 °C for 15 s, 63 °C for 20 s, 72 °C for 60 s). The cDNA of all samples was then amplified using 13 PCR cycles (Veriti 96-Well Thermal Cycler, Applied Biosystems, Thermo Fisher Scientific). After a cleanup (SPRIselect, Beckmann Coulter, Germany) as well as quality check and quantification (2100 Bioanalyzer, Agilent Technologies, Germany), the cDNA was sent for library construction and sequencing (Cologne Center for Genomics, University of Cologne, Germany). The libraries were sequenced on an Illumina Novaseq 6000 with a sequencing depth of about 100,000 reads/ spot covered by tissue. All raw sequence data were deposited in the SRA and are publicly available as of the date of publication.

### Mapping and quantification

Tissue and fiducial frames of each capture area were detected and aligned manually using the Visium Manual Alignment Wizard in Loupe Browser (v6.0.0) (10x Genomics). Sequencing data was then processed and mapped to a modified version of the *F. hepatica* genome (BioProject PRJNA179522, annotation version 2021-12-WormBase, WBPS17 + mitochondrial genome (Gene bank accession NC_002546.1)) using the spaceranger count pipeline (v1.3.1) (10x Genomics). The modified genome is available from the authors upon request. Approximately 63% out of the mean 118,604 reads per spot were confidently mapped to the transcriptome. Our four sections covered a total number of 2,020 spots, which captured a median of 2,192 genes and 6,138 UMI counts per spot.

### Data processing and clustering

Data from spaceranger was imported into Seurat (v4.3.0)^16,17^ for further processing. First, a quality control was carried out by looking at the total number of UMIs and genes as well as the percentage of mitochondrial genes per spot (Figure S1). No filters were applied, so that all tissue-covered spots were included in the further analysis. Data was normalized and scaled using SCTransform with default parameters for each sample individually. Datasets were then merged using Seurat. On the merged dataset we ran RunPCA for dimensionality reduction and RunHarmony (v0.1.1)^74^ for batch correction (dims.use = 1:8, theta = 0, lambda = 4.7). The number of principal components for dimensionality reduction was determined by visual inspection of the ElbowPlot provided by Seurat. We then ran FindNeighbors (reduction = "harmony", dims = 1:8), FindClusters (resolution = 3) and RunUMAP (reduction = "harmony", dims = 1:8) to receive a first clustering. The clusters were then inspected and annotated according to the underling tissue type (vitellarium, tegument, parenchyma, gut, uterus, testis, ovary and Mehlis’ gland). For a small number of spots, the cluster ID did not correspond to the tissue seen in the histological image. Therefore, we exported barcodes with corresponding cluster IDs & UMAP coordinates from Seurat to Loupe Browser to manually correct mismatching cluster IDs of those spots. New cluster IDs were then re-imported into Seurat to proceed with marker gene analysis.

### Identification of marker genes

We used the FindAllMarkers function provided by Seurat to identify marker genes for each cluster (test.use = ”roc”, only.pos = TRUE) (Table S2). Gene descriptions for all gene IDs were downloaded from WormBase ParaSite (BioMart) and then attached to the list of marker genes. Following the release of WBPS18, we adapted gene descriptions in our marker list according to this new version. Spatial expression patterns of marker genes were visualized using the SpatialFeaturePlot function in Seurat (crop = FALSE, pt.size.factor = 1, alpha = c(0.1,1), image.alpha = 0, stroke = 0.5,). Additionally, the DotPlot function was used to provide an overview of the expression of multiple genes.

To identify marker genes for the two cell types of the Mehlis’ gland, we manually selected spots covered by S1 and S2 cells in the Loupe Browser (Figure S3.2). Spots that were covered by other organs and tissues were labeled as "other." Barcodes and corresponding cluster IDs were then imported into Seurat to perform differential gene expression analysis. We used the FindMarkers function to identify marker genes for each celltype (test.use = ”wilcox”, assay = “SCT”, only.pos = TRUE) (Table S3). Annotations were added as described above.

### Gene ontology enrichment analysis

Gene Ontology (GO) annotation for *F. hepatica* was obtained from WormBase ParaSite (PRJNA179522, WBPS17) and added on by running InterProScan (v5.60.92.19)^75^. GO term enrichment analysis was performed using topGO (v2.46.0)^76^. Only marker genes with power values above the 25% percentile of each cluster were included in this analysis (“top 75%”). Analysis was done with weight01 method for all categories (BP and MF) with a node size restricted to ≥10. Significance was determined using Fisher’s exact test against all expressed genes.

### STRINGdb analysis

Molecular interactions were predicted using the STRING online tool (v11.5)^77^ after uploading the *F. hepatica* proteome (PRJNA179522). Gen-IDs for the top 75% of marker genes of each cluster were retrieved from our marker list and entered as a multiple protein search. Default settings were used to predict interactions with a minimum interaction (confidence) score of 0.4 (medium level of confidence).

### *In silico* investigation of *F. hepatica* gene families

We used the keyword search in WormBase ParaSite to identify members of the *F. hepatica* β-tubulin, PKC, glutathione S-transferase and ABC transporter families within the *F. hepatica* genome (PRJNA179522). To compare and assign those sequences to known liver fluke or human sequences provided in NCBI, we used reciprocal BLAST searches with standard parameters. In addition, SMART^78^ analysis was performed to confirm the domain structure of selected isoforms. *F. hepatica* orthologues of *F. gigantica* Ly6 proteins, recently characterized by Davey et al.^31^, were identified by WormBase ParaSite BLASTp with standard parameters. Interspecies orthologues were defined as Ly6 proteins with >85% amino acid identity.

Alignments of tubulin and PKC-β catalytic domain sequences were produced in Clustal Omega (v1.2.4)^79^. The tubulin alignment was annotated in Jalview (v2.11.2.7)^80^. For phylogenetic tree construction, β-tubulin, GST and ABC transporter sequences were aligned using the MUSCLE alignment provided within the Molecular Evolutionary Genetics Analysis (MEGA, v10.2.4) software version X^81^. Phylogenetic trees were then constructed by the maximum likelihood method and JTT matrix-based model with 1000 bootstrap replicates using MEGA X.

To explore the spatial expression of all family members, spatial plots were generated as described above. For better comparison of gene expression levels, we created a uniform color scale for some subsets of genes. Heatmaps were generated using the pheatmap package (v1.0.12)^82^ after calculating the average expression of each gene per cluster. Gene expression values were centered and scaled for each gene individually.

### Riboprobe synthesis

Templates for riboprobe synthesis were generated by TA-mediated cloning of pre-amplified cDNA sequences (Q5 High-Fidelity DNA Polymerase, New England Biolabs, Germany & AccuPrime Taq DNA Polymerase, High Fidelity, Thermo Fisher Scientific) with an average length of 400-500 bp. Primers for all genes can be found in Table S6. The resulting PCR product was ligated (T4 Ligase, New England Biolabs) with *AhdI*-digested (New England Biolabs) pJC53.2^83^ and used to transform NEB® 10-beta competent *E. coli* (New England Biolabs). Cloned sequences were confirmed by Sanger sequencing (Microsynth Seqlab, Germany) (sequencing primer: 5’-TTCTGCGGACTGGCTTTCTAC-3’ ^84^) and WormBase ParaSite BLAST (see Table S6 for results). Templates for *in vitro* transcription were generated by PCR amplification (Q5 High-Fidelity DNA Polymerase, New England Biolabs) from plasmids using a T7 extended primer (5’-CCTAATACGACTCACTATAGGGAG-3’ ^85^). *In vitro* transcription was finally performed using T3 or SP6 RNA polymerases (Roche, Germany) with the addition of Digoxigenin-11-UTP (Roche).

### *In situ* hybridization

*In situ* hybridizations (ISH) (chromogenic (CISH) or fluorescent (FISH)) were performed as described earlier with slight modifications^86^. 10 µm cryosections from adult *F. hepatica* were prepared as described above, post-fixed in 4% formaldehyde/PBS and permeabilized with PBSTx (0.5% Triton X-100). To inactivate endogenous peroxidase activity, slides were incubated in 0.03% H2O2/4x saline sodium citrate buffer (SSC) before continuing with prehybridization (FISH only). Hybridization reaction was carried out at 55 °C overnight. Probes were used at 1 µg/ml in hybridization buffer. The next day, multiple washing steps with hybridization washing buffer and decreasing concentrations of SSC were carried out, followed by blocking and antibody incubation (anti-DIG-POD (FISH) or anti-DIG-AP (CISH), Roche). After washing with maleic acid buffer (+ 0.1% Tween-20), the development reaction was carried out using the TSA Plus Cyanine 3 Kit (Akoya Biosciences, US) (FISH) or BCIP/NBT (Roche) in alkaline phosphatase buffer (CISH). For FISH, tissue sections were counterstained with Hoechst 33342 (1 µg/ml) or DAPI (0.1 µg/ml) and mounted with ROTImount FluoCare (Carl Roth, Germany). For CISH, 80% glycerol was used for mounting.

Imaging of *in situ* hybridizations was performed on an Olympus IX81 microscope (Olympus, Japan) or a Leica DM IL microscope (Leica Microsystems). Fiji (ImageJ, v1.54f)^87^ was used for linear adjustment of brightness and contrast of acquired images. Numbers of ISH experiments performed for each gene are listed in Table S6.

### In vitro culture and inhibitor treatment

The anthelminthic activity of the PKCβ inhibitor ruboxistaurin (LY333531) against adult stages of *F. hepatica* was assessed *in vitro* using different inhibitor concentrations (20, 50, or 100 μM). Worms were obtained as described above and cultured in RPMI medium (supplemented with 5% chicken serum and 1% ABAM-solution [10,000 units penicillin, 10 mg streptomycin, and 25 mg amphotericin B per ml], all from Gibco) at 37 °C in a 5% CO2 atmosphere for 72 h. Triclabendazole (20, 50, or 100 μM) served as positive control, the solvent dimethyl sulfoxide (DMSO) equivalent to the highest drug concentration as negative control. Medium plus inhibitor was refreshed every 24 h. Inhibitor-induced effects on worm viability were assessed using a stereo microscope at 10× magnification (M125 C, Leica, Germany) and the following scores: 3 (normal motility), 2 (reduced motility), 1 (minimal and sporadic movements), and 0 (dead).

## Acknowledgements

Financial support by the Deutsche Forschungsgemeinschaft (DFG) under grant HA 6963/2-1 and by the State of Hesse, LOEWE Center "DRUID", is gratefully acknowledged. O.P. received a scholarship by the Justus Liebig University Giessen. We would also like to thank Dr. Knut Beuerlein at the Rudolf Buchheim Institute of Pharmacology, JLU, for providing and supporting us with their Leica DMi8 microscope. We also thank the group of Prof. Bernhard Spengler (Institute of Inorganic and Analytical Chemistry, JLU) for access to their cryostat.

## Author contributions

Conceptualization, S.G., S.H.; Methodology, S.G., O.P.; Investigation, S.G., O.P., T.S.; Visualization, S.G.; Resources, Z.L.; Writing – Original Draft Preparation, S.G.; Writing – Review & Editing, all authors; Supervision and Funding Acquisition, S.H.

## Declaration of interests

The authors declare no conflict of interest. The funders had no role in study design, data collection and analysis, decision to publish, or preparation of the manuscript.

## Supplementary figures and legends

**Figure S1.**
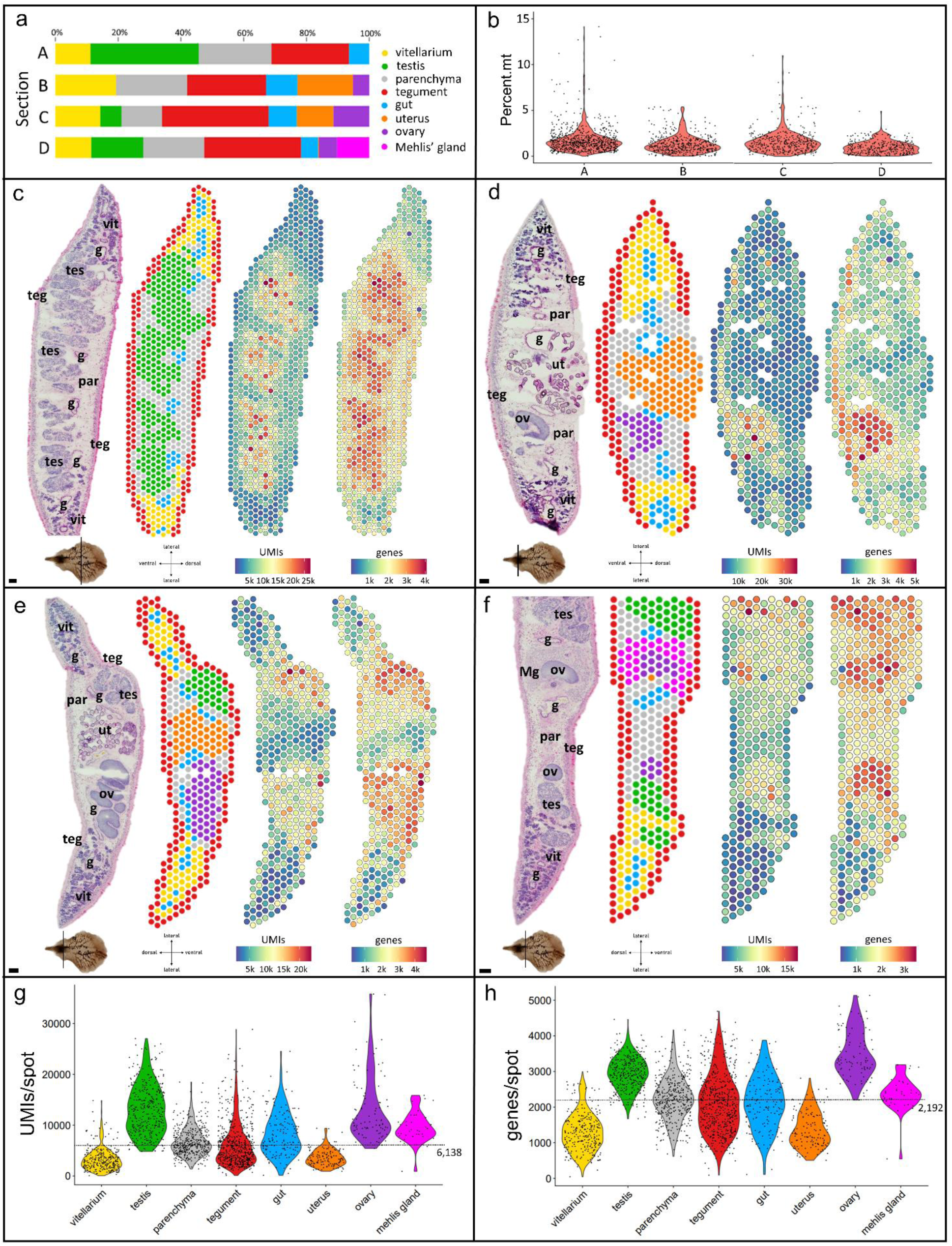
Tissue composition and QC metrics of all four specimens in the liver fluke spatial transcriptomics dataset. **(a)** Tissue (cluster) composition of sections A-D. **(b)** Violin Plot showing the percentage of mitochondrial genes/ spot split by section. **(c-f)** Left to right: H&E stained tissue sections and corresponding spatial projections of 777, 481, 412 and 350 mRNA-binding spots covered by these tissue sections, respectively. Sectioning plane and orientation are indicated underneath. Scale = 100 µm, g: gut, Mg: Mehlis’ gland, ov: ovary, par: parenchyma, teg: tegument, tes: testis, ut: uterus, v: vitellarium. The first spatial projection shows the cluster assignment of all spots. Clusters are colored according to (a) and Figure 1. The second and third projection are displaying the distribution of UMI (Unique Molecular Identifier) and gene counts, respectively. The number of UMIs/ genes is encoded by color (blue – red = low – high). **(g,h)** Violin plot displaying UMI/ gene counts per spot, split by cluster. The median UMI/gene count across all spots is indicated by the dashed line.

**Figure S2:**
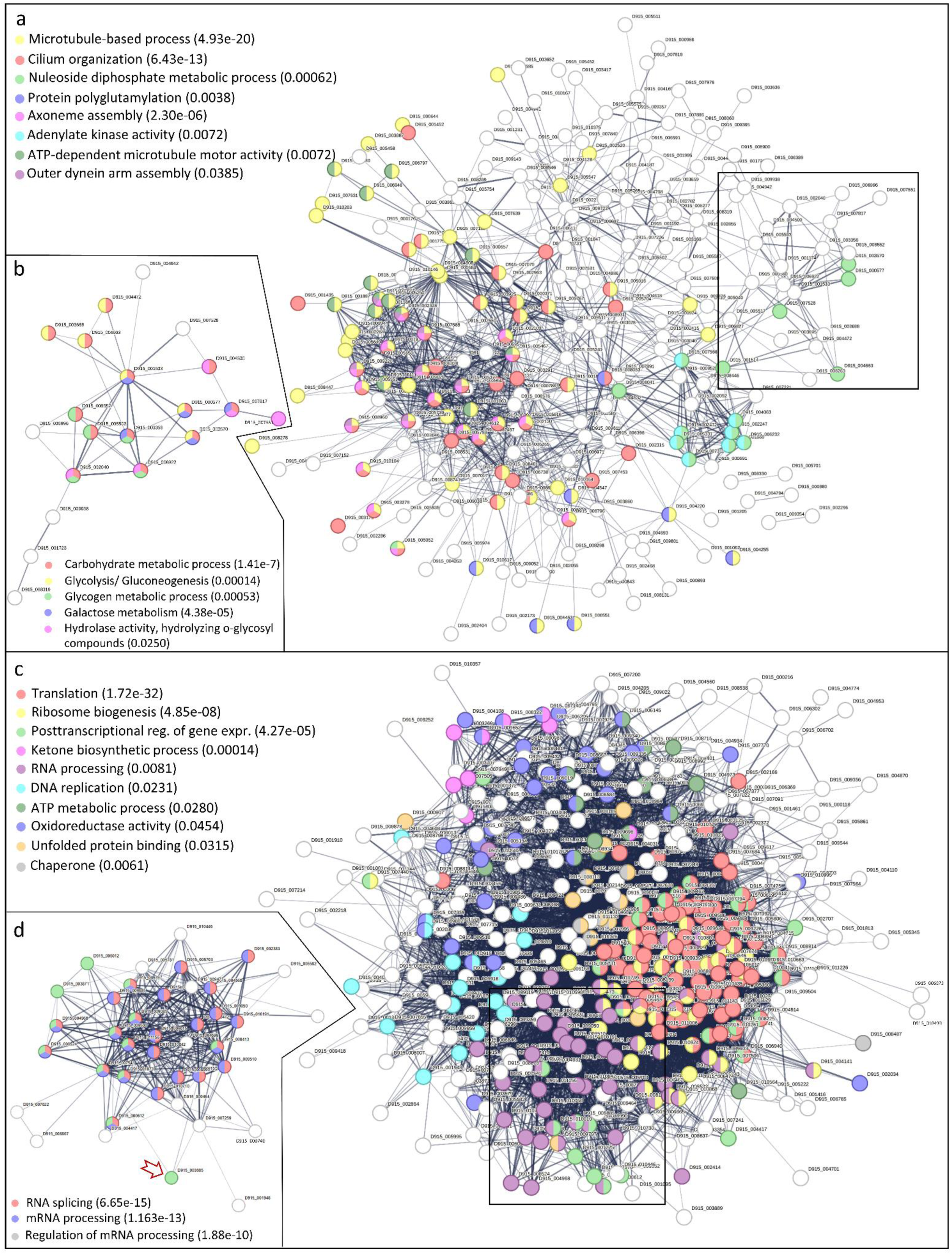
STRING analyses for marker genes of ovary and testis. STRING analysis of marker genes (top 75%) for **(a)** testis and **(c)** ovary. Default settings were used to predict interactions with a minimum interaction (confidence) score of 0.4 (medium level of confidence). Lines (edges) are indicating interactions that were predicted based on the function of homologues. Disconnected nodes are not shown. Functional enrichments in the network were predicted by STRING. Selected terms are highlighted in color and labelled (false discovery rate in brackets). Subnetworks in **(b)** and **(c)** resulted from kmeans clustering in STRING (number of clusters = 10). Rectangles in (a) & (b) mark where the corresponding genes are located in the full network. **(d)** Arrow indicates Zinc finger CCCH domain-containing protein 3 (D915_003685).

**Figure S3.1.**
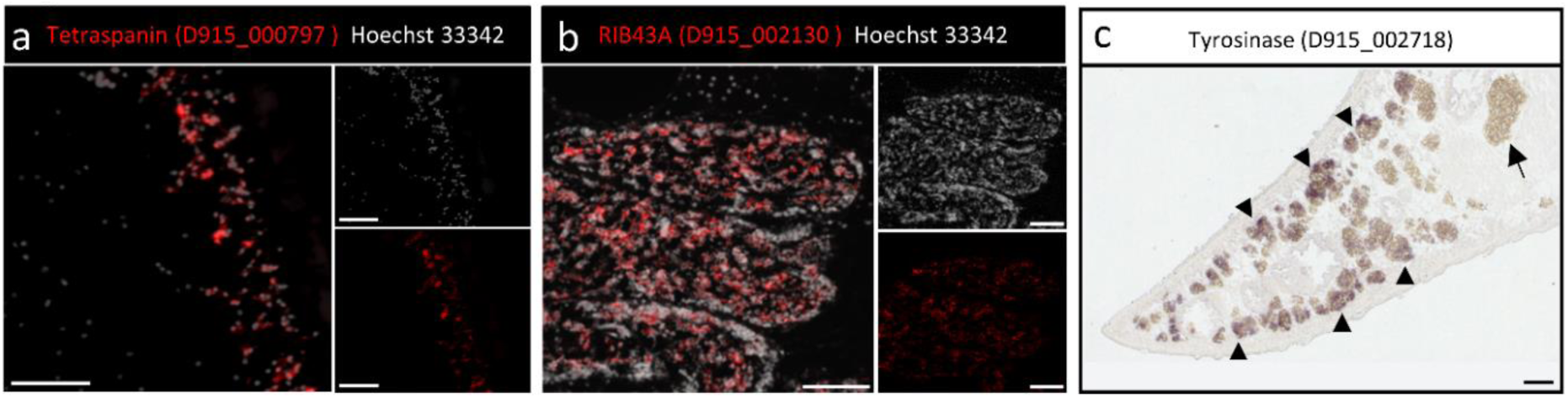
(F)ISH validation of additional marker genes. **(a)** Fluorescent in situ hybridization (FISH) of the tegument marker Tetraspanin (D915_0000229). **(b)** FISH of the testis marker RIB43A (D915_002130). **(c)** Chromogenic ISH of the vitellarium specific tyrosinase (D915_002718). Black arrowheads indicate indigo dye deposition within vitelline follicles. No staining was observed within mature vitelline cells in the vitelline duct (black arrow). (a-c) Scale = 100 µm.

**Figure S3.2.**
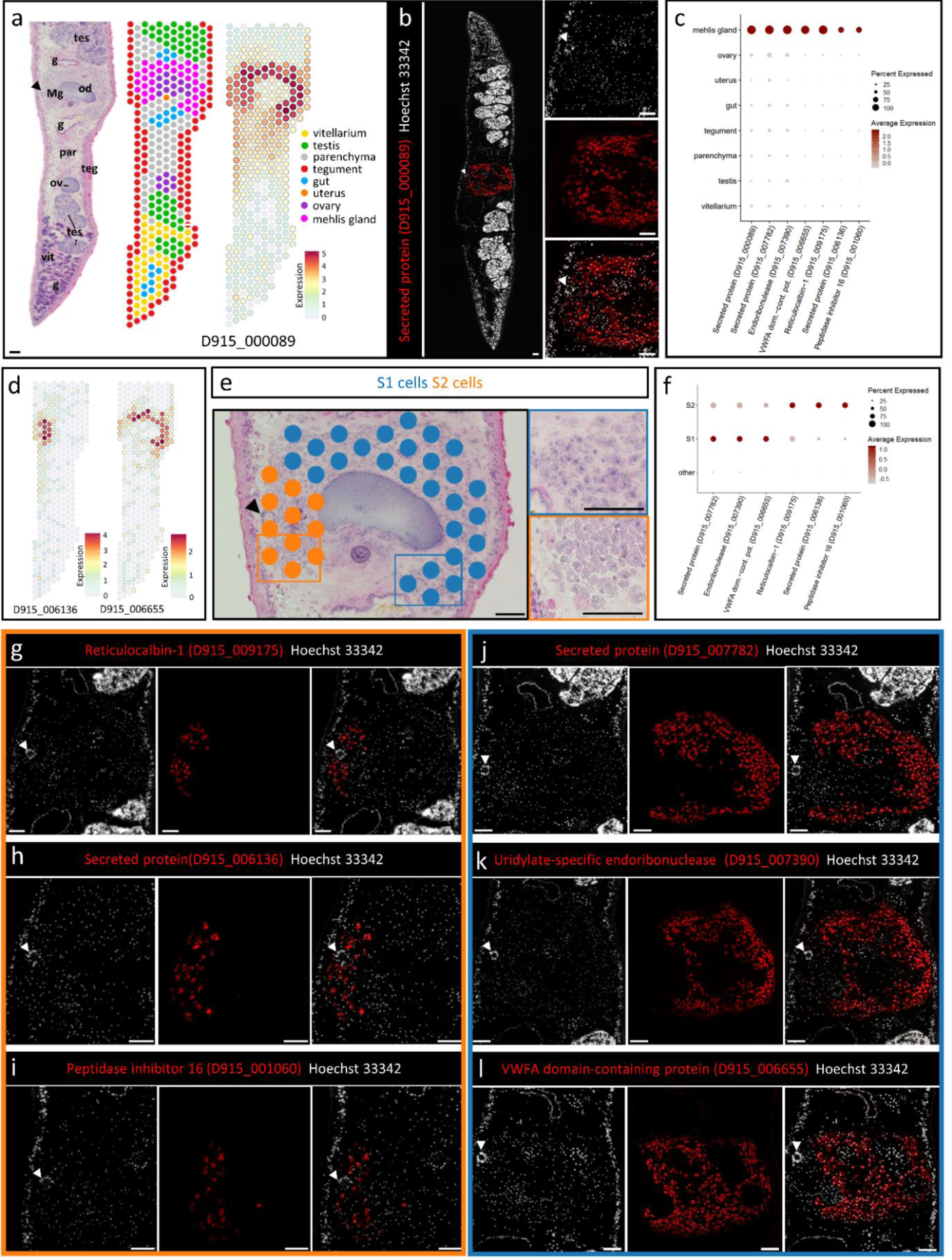
The liver flukes Mehlis’ gland is composed of two transcriptionally distinct cell types. **(a)** Mehlis gland tissue was only included in one of our four specimens (“section D”). Left to right: H&E stained tissue sections and corresponding spatial projections of 350 mRNA-binding spots covered by this tissue section. g: gut, Mg: Mehlis’ gland, ov: ovary, par: parenchyma, teg: tegument, tes: testis, v: vitellarium. The first spatial projection shows the cluster assignment of all spots. Clusters are colored and labelled. The second spatial projection shows the spatial expression of the Mehlis’ gland marker D915_000089. This secreted protein was tissue-wide expressed within the Mehlis’ gland. Expression level encoded by color (grey = low, red = high). **(b)** D915_000089 expression was confirmed by fluorescent in situ hybridization (FISH). Overview (left) and detail (right). **(c)** DotPlot showing expression profiles of selected tissue markers of the Mehlis’ gland cluster. Dot color encodes the average expression level (scaled) across all spots within a cluster. Dot size encodes the percentage of spots within a cluster that have captured this transcript. Please note: While spatial plots (a,d) are shown for only one representative section, the DotPlot includes expression data from all four tissue sections in the data set. **(d)** Several marker genes of the Mehlis’ gland cluster, e.g. another secreted protein (D915_006136) and a VWFA domain containing protein (D915_006655), showed a more localized expression within either the dorsal or the lateral and ventral part of the gland. Expression level encoded by color (grey = low, red = high). **(e)** These locations correspond to the arrangement of S1 (blue) and S2 cells (orange) within the Mehlis’ gland of *F. hepatica* [S1]. Having a closer look at the histology of the tissue section, we were also able to detect distinct histological features described for those cell types: S1 cells are smaller and have a denser and basophilic cytoplasm compared to S2 cells which are larger and have a paler and more eosinophilic cytoplasm [S1].Therefore, we were able to manually select spots covering either S1 or S2 cells using Loupe Browser (10x Genomics). **(f)** We then performed differential gene expression analysis in Seurat to compare the gene expression of both cell types. DotPlot showing expression profiles of selected tissue markers of Mehlis’ gland S1 & S2 cells. “Other” includes all remaining spots in the spatial dataset (all remaining clusters/tissues in all four sections). Dot color encodes the average expression level (scaled) across all spots within a cluster. Dot size encodes the percentage of spots within a cluster that have captured this transcript. For a complete list of cell type markers see Table S2. **(g-i)** We found and confirmed by FISH that S1 cells were characterized by the expression of Poly(U)-endoribonuclease (D915_007390), a von Willebrand factor A (VWFA) domain-containing protein (D915_006655), and a C3H1-type domain-containing protein (D915_007782). **(j-l)** For S2 cells we identified reticulocalbin (D915_009175), peptidase inhibitor 16 (D915_001060) and another secreted protein (D915_006136) to be exclusively expressed within this cell type. (a,b,e,g-l) Scale = 100 µm, arrowhead marks Laurers’ channel entering Mehlis’ gland from dorsal.

**Figure S4.**
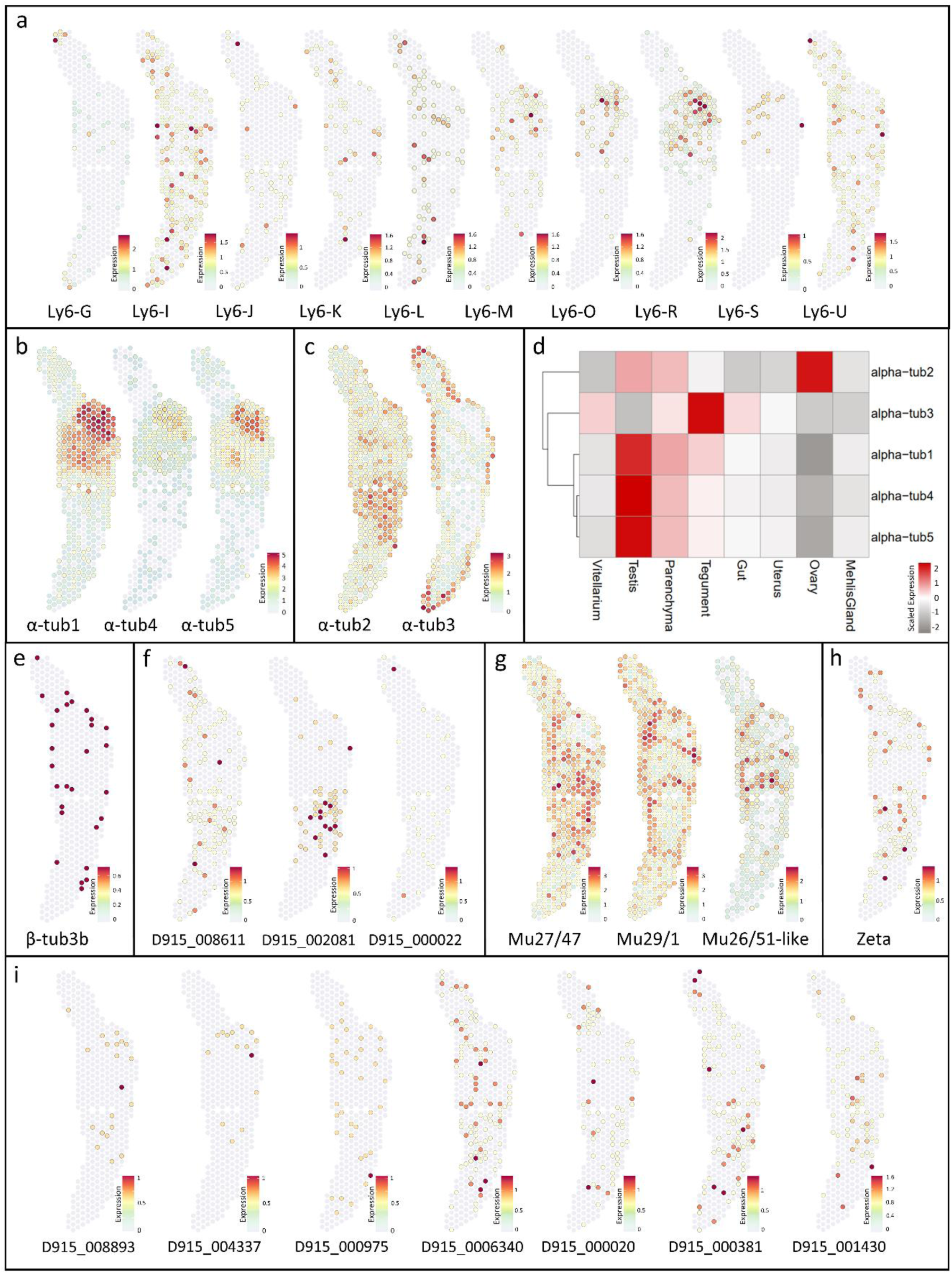
Spatial expression patterns of additional Ly6 proteins, tubulins, PKCs, GSTs and ABCB transporters. **(a)** Spatial projections showing expression patterns of *F. hepatica* Ly6 proteins not shown in Figure 3: Ly6-G (D915_008997), Ly6-I (D915_002959), Ly6-J (D915_006710), Ly6-K (D915_009743), Ly6-L (D915_008235), Ly6-M (D915_008952), Ly6-O (D915_000989), Ly6-R (D915_000988), Ly6-S (D915_000991), Ly6-U. **(b,c)** Spatial projections showing expression patterns of *F. hepatica* α-tubulins. α-tubulin isoforms with predominant expression in the testis: α-tub1 (D915_009242), α-tub4 (D915_009559), α-tub5 (D915_005959). **(c)** α-tub2 (D915_005370), α-tub3 (D915_8616). **(d)** Heatmap showing the average α-tubulin isoform expression per cluster. Expression values were centered and scaled for each row (each gene) individually. Please note: While spatial plots (b,c) are shown for only one representative section, the Heatmap includes expression data from all four tissue sections in the data set. **(e)** Spatial projection showing expression patterns of *F. hepatica* β-tubulin 3b (“beta-tub3”/ D915_002077). **(f)** Spatial projections showing expression patterns of PKCs not shown in Figure 5. **(g,h)** Spatial projections showing expression patterns of glutathione S-transferases not shown in Figure 6. mu class GSTs: Mu27/47 (D915_010266), Mu29/1 (D915_007977), Mu28/51-like (D915_008366). zeta class GST (D915_006391). **(i)** Spatial projections showing expression patterns of ABC transporters (subfamily B) not shown in Figure 6. (a-c & h-i) Expression level encoded by color (grey = low, red = high).

**Figure S5.**
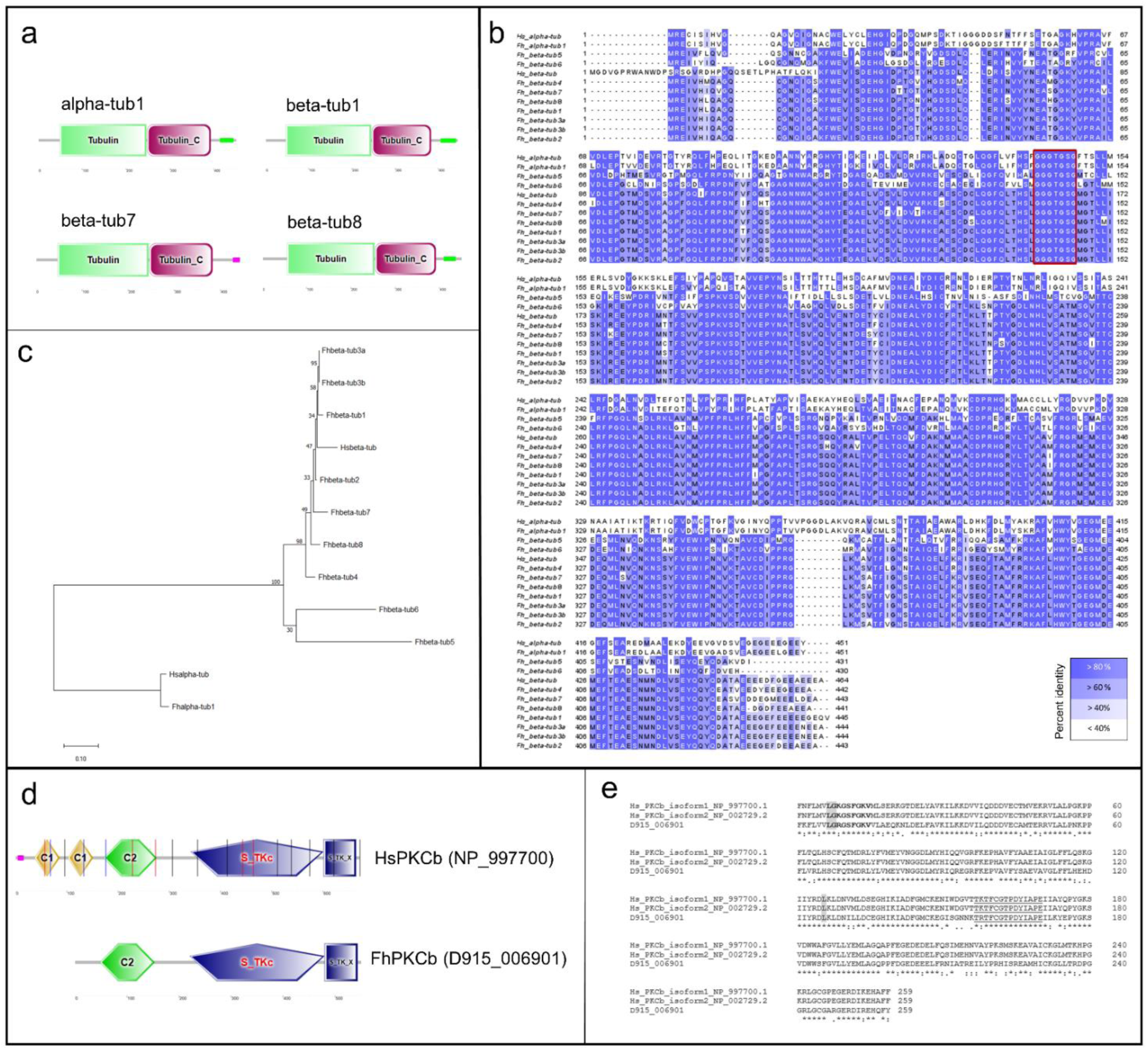
*In silico* characterization of β-tubulins and PKCs. Related to Figure 5 and STAR Methods. **(a-c)** The keyword search in WormBase ParaSite was used to identify members of the *F. hepatica* β- tubulin family within the *F. hepatica* genome (PRJNA179522). In addition to sequences of the known β-tubulin isotypes 1-6, we identified two previously undescribed β-tubulins (β-tub7 (D915_000542) and β-tub8 (D915_000946)). β-tub1 (D915_007398), β-tub2 (D915_002311), β-tub3a (D915_004911), β-tub3b (D915_002077), β-tub4 (D915_001342), β-tub6 (D915_008457). β-tubulin isoform 5 was found fragmented with D915_005076 and D915_003963 representing two halves of the complete ß- tubulin isoform 5. Therefore, these sequences were concatenated and used as the *F. hepatica* isoform 5 gene for alignment and phylogenetic tree construction. **(a)** SMART analysis confirmed the tubulin domain structure of *F. hepatica* β-tub7 and β-tub8 in comparison with α-tub1 (D915_009242) and β- tub1. **(b,c)** β-tub7 and β-tub8 had 90-95% amino acid identity with *F. hepatica* beta tubulin isoforms 1-4, but only 41% with alpha-tub1. **(b)** Alignment was created with Clustal Omega and colored using the PID (percent identity) option in Jalview. In addition to all *F. hepatica* (Fh) β-tubulin sequences, we included the amino acid sequence of α-tub1 and one human (Hs) alpha (NP_006000.2) and beta tubulin (NP_110400.1) sequence each. The tubulin signature sequence (GGGTGSG) is marked (red rectangle). Maximum likelihood analysis for phylogenetic tree construction was performed in MegaX to 1000 bootstraps with JTT substitution. α-tubulin sequences served as outgroup. **(d,e)** D915_006901 was confirmed as PKCβ orthologue by BLAST search starting from the human PKCß amino acid sequence (NP_997700.1) and re-blast against *H. sapiens*. **(d)** SMART analysis of human and liver fluke PKCβ. The domain composition (C2 regulatory domain in front of the kinase domain) classifies both as conventional PKCs [S2]. **(e)** Alignment of PKCβ catalytic domains. The catalytic domain of human PKCβ, which is bound by ruboxistaurin, shows 73.36% identity to the *Fasciola* orthologue. The conserved ATP binding site (**bold**) and the activation loop residues (<u>underlined</u>) are marked. The main residues involved in ruboxistaurin binding obtained for the human sequence [S2] are Lys349, Gly350 and Lys467 (grey).

**Figure S6.**
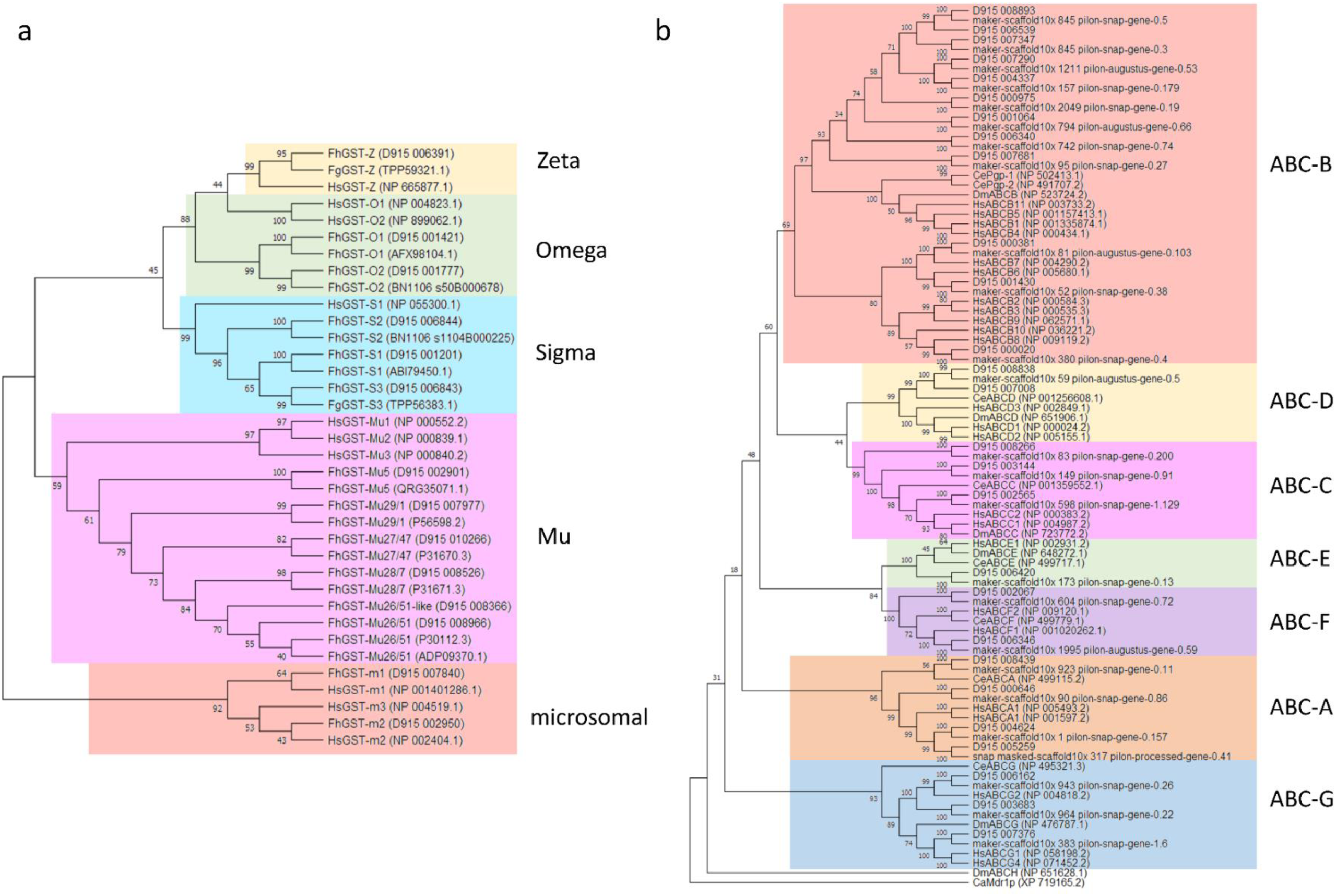
Class/ Subfamily assignment of GSTs and ABC transporters by phylogenetic tree construction. Phylogenetic analysis of *F. hepatica* (Fh) glutathione S-transferases (GSTs). GST amino acid sequences from the “St. genome (PRJNA179522, “D915”-IDs) were analyzed together with human (Hs) GST sequences as well as known *la* GST sequences. The latter were adopted from Stuart et al. [S3]. If no *F. hepatica* sequence was available, *F. ica* (Fg) sequences were used instead. Maximum likelihood analysis was performed in MegaX to 1000 bootstraps TT substitution. The tree was displayed as “Topology only”. Microsomal GST sequences were used as outgroup. **(b)** genetic analysis of *F. hepatica* ATP-binding cassette (ABC) transporters. ABC transporter amino acid sequences of t. Louis” *F. hepatica* genome (PRJNA179522, “D915”-IDs) and “Liverpool” genome (PRJEB25283, “marker-scaffold”- ere analyzed together with human (Hs), *C. elegans* (Ce) and *D. melanogaster* (Dm) ABC sequences of all subfamilies *Candida albicans* (Ca) Mdr1p was used as outgroup. Liverpool ABC transporter gene IDs were adopted from Beesley S4]. Maximum likelihood analysis was performed in MegaX to 1000 bootstraps with JTT substitution. The tree was yed as “Topology only”.

## Supplementary video and Excel table titles and legends

**Video S1. Adult *Fasciola hepatica* depicts normal motility (score 3) after 72 h treatment with DMSO as control**.

**Video S2. Adult *Fasciola hepatica* with a motility score of 0 (“dead”) after 72 h treatment with 50 µM ruboxistaurin**.

**Table S1. List of samples used for spatial transcriptomics and corresponding metrics**

**Table S2. List of marker genes per cluster**

The FindAllMarkers() function embedded in Seurat was used to identify markers for each of the clusters in the spatial transcriptomics dataset by “ROC” test.

**Table S3. Differential gene expression of Mehlis’ gland S1 and S2 cells**

The FindMarkers() function embedded in Seurat was used to identify upregulated genes in S1 vs S2 cells and vice versa by “wilcox” test.

**Table S4. Summary of all FhLy6 proteins**

**Table S5. Reagents & chemicals used within the Visium Spatial Gene Expression workflow in deviation from manufacturer’s instructions**

**Table S6. Riboprobes for *in situ* hybridization**

Cloning information for all markers used for ISH validation and number of independent *in situ* hybridization experiments performed per marker gene.

## Notes

### Competing Interest Statement

The authors have declared no competing interest.

